# Chemoproteomic Characterization of GPX4 Covalent Ligands and Targeted Degradation

**DOI:** 10.64898/2026.04.29.721679

**Authors:** Vilas D. Kadam, Guangcan Bai, Chen Mozes, Hanling Guo, Zhouyiyuan Xue, Qi Miao, Jian Wang, Minhua Li, Feng Li, Daisuke Nakada, Zhi Tan, Xiaoyu Zhang, Mingxing Teng

## Abstract

Despite intensive efforts, the ferroptosis gatekeeper glutathione peroxidase 4 (GPX4) remains difficult to selectively target due to stringent structural constraints surrounding its catalytic selenocysteine, which impose tight requirements on warhead reactivity and geometry. Here, leveraging a chemoproteomic approach, we characterize a potent and selective covalent GPX4 inhibitor featuring a pyrimidinylmethyl isourea warhead and define the chemical features underlying its proteome-wide selectivity. This chemotype enables tunable electrophile reactivity through steric and electronic modulation of leaving group ability, suggesting potential broader utility for targeting other recalcitrant proteins. Building on this scaffold, we further develop two selective GPX4 degraders - one CRBN-dependent and the other CRBN-independent - enabling complementary modulation of GPX4 through both inhibition and degradation. Together, these molecules expand the GPX4 chemical toolbox for more nuanced interrogation of GPX4 biology.

## INTRODUCTION

Ferroptosis is a regulated form of cell death driven by iron-dependent lipid peroxidation, with broad implications in diseases such as cancer, neurodegeneration, ischemia-reperfusion injury, infection, and sterile inflammation.^1-5^ GPX4 serves as a critical gatekeeper of ferroptosis by using glutathione (GSH) to reduce phospholipid hydroperoxides to their corresponding alcohols, thereby preventing oxidative lipid damage.^6,7^ Together with parallel defense pathways, including the ferroptosis suppressor protein 1 (FSP1)-ubiquinol (CoQH_2_) system, the dihydroorotate dehydrogenase (DHODH)-CoQH_2_ system, and the GTP cyclohydrolase 1 (GCH1)-tetrahydrobiopterin (BH_4_) system, the GPX4-GSH axis constitutes a core component of the cellular ferroptosis defense network,^3^ making GPX4 an attractive therapeutic target.^8^ Mechanistically, GPX4 catalyzes the reduction of phospholipid hydroperoxides through oxidation of its active-site nucleophilic selenol (Se-H) to selenenic acid (Se-OH), which is subsequently reduced by GSH via a selenenyl sulfide (Se-SG) intermediate to revert to the active selenol form. Mutation of this catalytic selenocysteine (Sec46, hereafter referred to as U46) to cysteine reduces enzymatic activity by roughly 90%, likely due to an unfavorable shift in pKa, and increases sensitivity to redox stress,^9^ underscoring its essential role in GPX4 function.

The catalytic selenocysteine U46 within the GPX4 active site is highly amenable to covalent modification, a feature that has spurred extensive chemistry efforts to develop targeted covalent inhibitors. These endeavors have yielded a broad array of chemical scaffolds, including RSL3 and ML162, which utilize an α-chloroacetamide warhead,^10,11^ and DPI17, which features a pyrimidinylmethyl chloride moiety.^10^ More recent innovations have introduced diverse electrophilic chemistries, such as the nitrile oxide of JKE-1777,^12^ the 2-ethynylthiazole-4-carboxamide of A1,^13^ and various other scaffolds incorporating propiolamide or similar reactive groups (Figure 1).^14-17^ Yet, despite this expanding toolkit, critical challenges remain in the development of effective GPX4-targeting inhibitors. A primary hurdle is the absence of a well-defined, deep active-site pocket in GPX4. Structural analysis of the mutant cytosolic form GPX4^C66S^ in complex with the ligand (*S*)-ML162 reveals a striking paradox: although the ligand binds within the active site, a substantial portion of its structure extends into the solvent-exposed region without making clear protein-ligand interactions,^18^ nonetheless, it still maintains potent cellular target binding. This observation supports the hypothesis that, in a cellular context, GPX4 may function as a membrane-associated complex. This state is thought to be mediated by GPX4’s cationic patch, which - as proposed by several studies, ^19-23^ - facilitates the formation of a functional membrane-bound state that provides a ‘true’ binding pocket to accommodate GPX4 ligands. Consequently, enzymatic assays using recombinant GPX4 likely fail to fully recapitulate the protein’s endogenous state. Another significant hurdle lies in the presence of seven additional cysteine residues in cytosolic GPX4 variant (C2, C10, C37, C66, C75, C107, and C148). This abundance of thiols imposes a stringent requirement for site-specific modification at U46, demanding finely tuned warhead reactivity to prevent both intra-protein off-site labeling and broader proteome-wide off-target effects. Furthermore, attenuating the intrinsic reactivity of covalent inhibitors is critical to reduce plasma clearance and enhance *in vivo* pharmacokinetics.^24^ Current gold-standard inhibitors, such as RSL3 and ML162, have been shown to label C66 indiscriminately,^25^ highlighting their lack of site-specificity. These limitations underscore the need for a novel class of electrophiles capable of selectively engaging U46 with moderate, controlled reactivity. Such an approach would bridge a critical gap in the chemical biology toolbox to enable effective *in vivo* applications. To date, to the best of our knowledge, no GPX4-targeting probe has successfully reconciled high cellular potency with rigorous proteome-wide selectivity.

**Figure 1.**
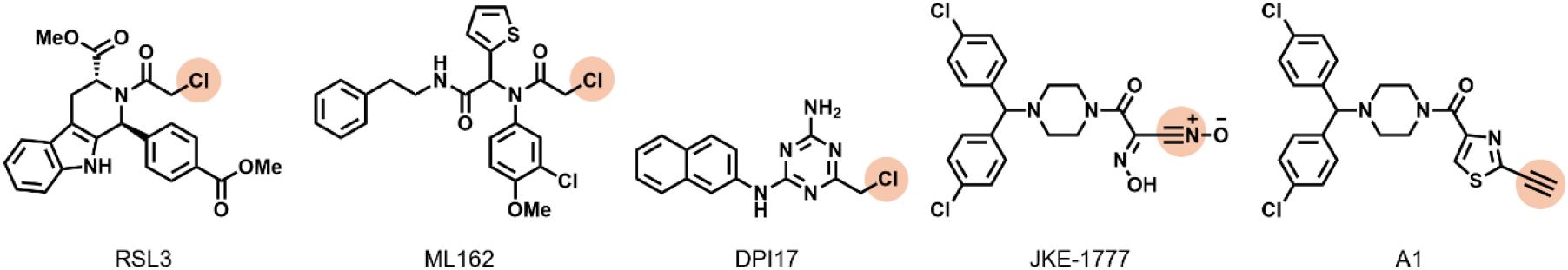
Chemical structures of representative GPX4 inhibitors. Warhead leaving groups or addition sites are highlighted with orange circles.

## RESULTS

A recent patent filing describes an intriguing molecule, referred to here as VK-3-189 (Ex. E46 in the patent),^26^ which features a novel pyrimidinylmethyl isourea motif designed to engage U46 through nucleophilic attack, functioning as a tailored leaving group. We synthesized VK-3-189 along with its enantiomer VK-3-188, the unsubstituted analog VK-3-200, and the pyrimidinylmethyl chloride VK-3-197 to roughly evaluate the impact of warhead stereochemistry, substitution, and warhead type on activity and reactivity (Figure 2A). The superior cell-killing effect of VK-3-189 compared with the reference benchmark RSL-3 and its analogs was confirmed in MDA-MB-231 breast cancer cells using a CellTiter-Glo^®^ luminescent viability assay. In combination with a rescue assay using the radical-trapping antioxidant ferrostatin-1 (fer-1),^27^ VK-3-189 exhibited a clear selectivity window between the two assays, suggesting it functions as a ferroptosis inducer (Figure 2B,C). Additionally, while warhead stereochemistry does not significantly affect kinetic solubility, it extends the half-life in both mouse and human liver microsomes (Figure 2C). Collectively, these results highlight the critical contribution of warhead architecture to both the cellular potency and favorable metabolic stability properties of VK-3-189. Next, London’s thiol reactivity assay,^28^ based on Ellman’s reagent (DTNB), was applied as a readily implementable method to assess the reactivity of VK-3-189 and its analogs. The assay involves incubating test compounds with reduced DTNB and monitoring the decrease in TNB^2-^ absorbance at 412 nm wavelength over 4 h. The alkylation rate constant (k) was determined by linear regression using a second-order kinetic model, with larger k values indicating faster reaction kinetics and higher intrinsic electrophile reactivity toward thiols. Accordingly, the pyrimidinylmethyl chloride VK-3-197 is approximately 5-fold more reactive than VK-3-189, whereas VK-3-189 exhibits similar kinetics to its enantiomer VK-3-188. Notably, the CF_3_ substitution-free analog VK-3-200 is roughly 1.4-fold less reactive than VK-3-189 (Figure 3). From a chemical perspective, this finding highlights the tunability of the pyrimidinylmethyl isourea warhead, where substituent steric and electronic effects can be leveraged to dial up or down leaving group ability and, consequently, fine-tune electrophile reactivity for context-dependent applications.

**Figure 2.**
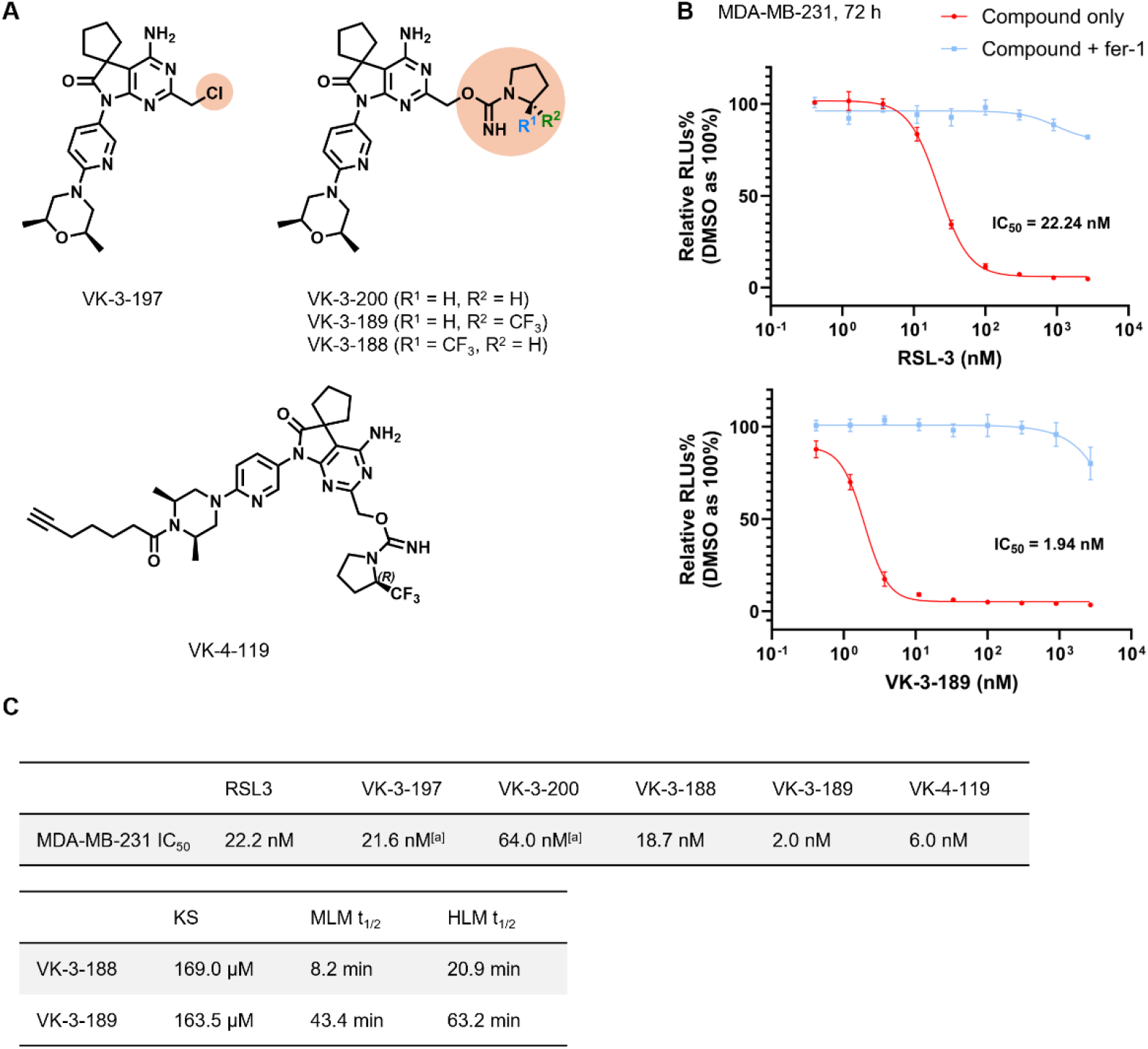
VK-3-189 is a potent cellular GPX4 inhibitor. (A) Chemical structures of VK-3-189, VK-3-188, VK-3-200, VK-3-197, and the alkyne-containing analog VK-4-119. Warhead leaving groups are highlighted with orange circles. (B) Cell-killing effects following 72 h treatment with VK-3-189 and the benchmark RSL3 in MDA-MB-231 cells, with rescue by fer-1 (1.5 μM). Data are shown as mean ± s.d. (n = 3 biological replicates). (C) Summary of 72 h cell-killing IC_50_ values for RSL3, VK-3-189, and related analogs, along with kinetic solubility (KS) and mouse/human liver microsome stability (MLM/HLM t_1/2_) for VK-3-189 and VK-3-188. [a] Data were normalized across batches using VK-3-189 as a reference.

**Figure 3.**
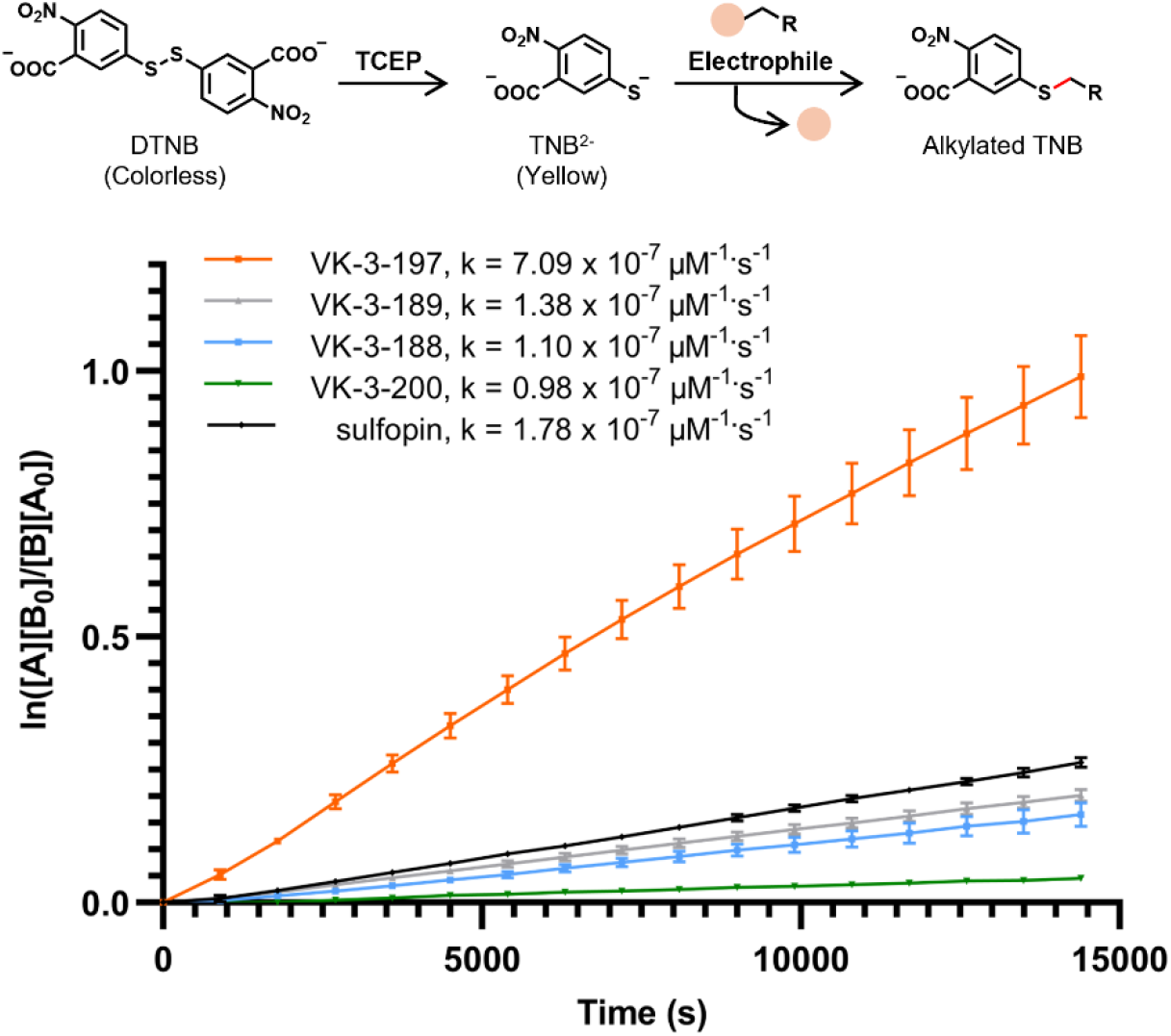
Thiol reactivity of VK-3-189 and analogs with distinct electrophile warheads. Thiol reactivity was assessed using London’s thiol reactivity assay. In the presence of TCEP, DTNB is reduced to TNB^2−^, which exhibits strong absorbance at 412 nm. Alkylation of TNB^2−^ by electrophilic fragments decreases this absorbance. Reaction kinetics were fitted to a second-order model, where [A] represents the electrophile concentration and [B] the TNB^2−^ concentration. Rates were determined by linear regression over 4 h, with each data point representing the mean of technical triplicates (n = 3). Due to the poor solubility of RSL3, the PIN1 inhibitor sulfopin - which shares RSL3’s α-chloroacetamide warhead - was used as a control.

We next sought to characterize proteome-wide selectivity of VK-3-189, initially employing cysteine-directed activity-based protein profiling (ABPP).^29-33^ Specifically, live cells were treated with VK-3-189 in the presence of the ferroptosis inhibitor liproxstatin-1 (lip-1) to ensure that the measured cysteine reactivity reflected direct ligand engagement rather than secondary effects of cell death. Following treatment and lysis, reactive cysteine sites were labeled with a desthiobiotin iodoacetamide (DBIA) probe. After trypsin digestion, probe-labeled peptides were enriched and analyzed via TMT-based quantitative proteomics. This site-specific competition assay enabled monitoring of VK-3-189 engagement across thousands of cysteine residues. The results revealed a highly targeted profile, with minimal off-target reactivity observed across the proteome, except for C282 in HMOX2. Notably, the GPX4 U46-containing peptide was not detected. This was likely due to the low abundance of selenocysteine combined with current sensitivity limits and the inherent challenges of mass spectrometry for selenium-containing peptides, such as poor ionization and complex fragmentation.^34-36^ However, other GPX4 residues, such as C66, C75, and C107, were successfully covered and showed no significant competition (Figure 4A, Data S1). This binding profile distinguishes VK-3-189 from previously reported GPX4 inhibitors such as RSL3, which often exhibit non-specific cysteine engagement on the GPX4 surface as well as other off-target proteins.^10,25,37^ Collectively, these data underscore the high proteome-wide selectivity of VK-3-189 at the residue level.

**Figure 4.**
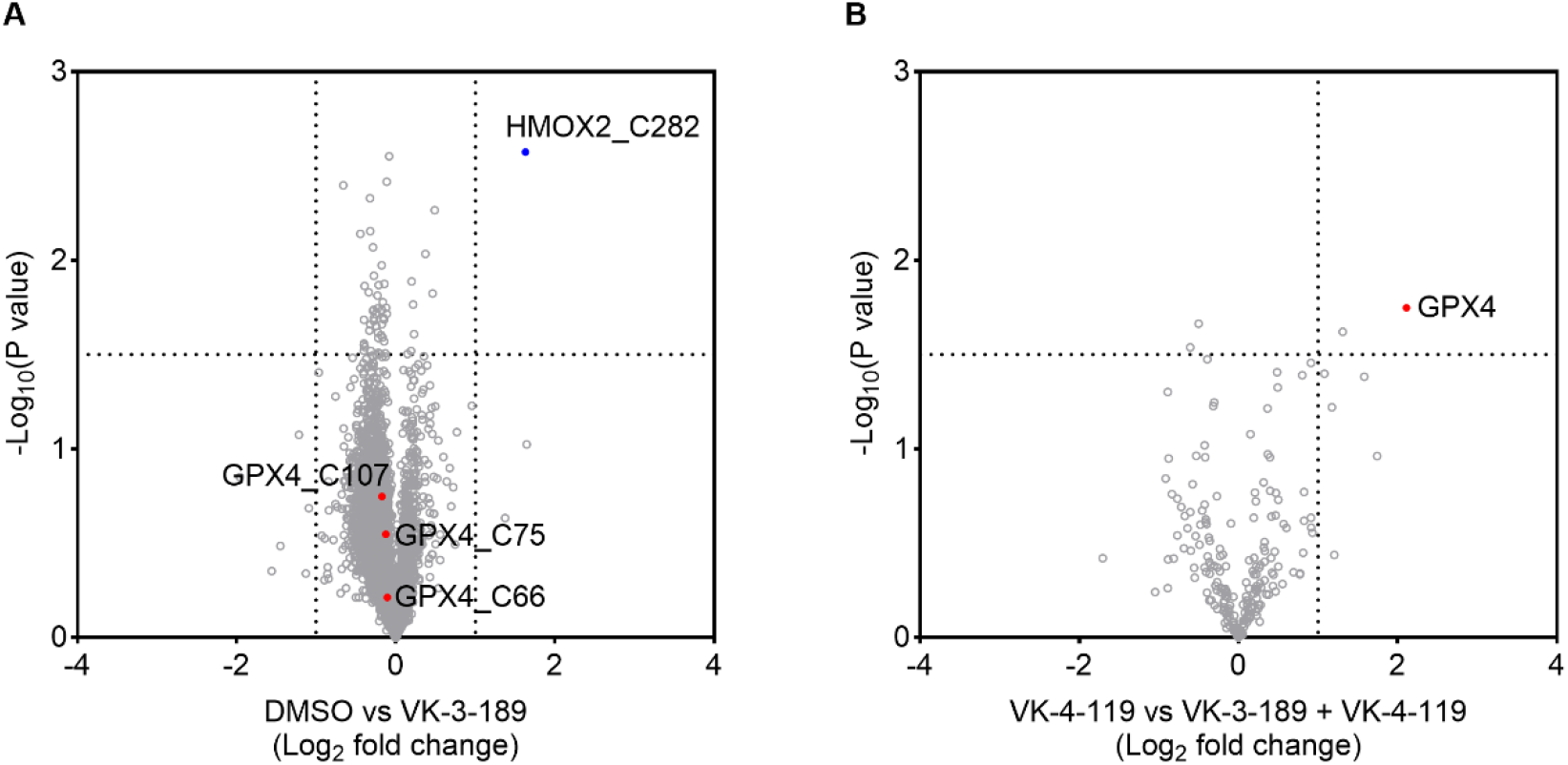
ABPP reveals proteome-wide selectivity of VK-3-189. (A) Cysteine-directed ABPP in HEK293T cells treated with DMSO or VK-3-189 (1 μM, 2 h), in the presence of lip-1 (2 μM) to prevent cell death. The GPX4 U46-containing peptide was not detected. (B) Protein-directed ABPP in HEK293T cells treated with VK-4-119 (1 μM, 2 h) alone or following co-treatment with VK-3-189 (3 μM), in the presence of lip-1 (2 μM).

To further validate the direct engagement of VK-3-189 with GPX4 at the protein level, we prepared an alkyne-functionalized analog, VK-4-119 (Figure 2A,C), to serve as a clickable capture probe for competitive ABPP.^12,38^ Briefly, cells were co-treated with VK-4-119 and lip-1 in the presence or absence of VK-3-189 for 2 h. This co-treatment format was chosen to ensure a direct assessment of target engagement while minimizing confounding cellular stress-responses that are not fully suppressed by lip-1. Following cell lysis, the cleared lysates were conjugated to a desthiobiotin-PEG3-azide linker via copper-catalyzed azide-alkyne cycloaddition (CuAAC).^39^ The labeled proteins were then precipitated, solubilized in urea, and subjected to reduction and alkylation. After enrichment on streptavidin beads, the proteome was digested on-bead using a Trypsin/LysC mixture. The resulting peptides were subsequently analyzed by TMT-based quantitative proteomics. Our results revealed a remarkably clean interaction profile: among the eight proteins significantly enriched by the alkyne probe (applying a Log_2_ fold-change cutoff of 2), only GPX4 displayed a substantial reduction in enrichment upon competition with VK-3-189 (Figure 4B, Data S1). This protein-level evidence complements our site-specific data, collectively confirming that VK-3-189 is a highly selective covalent GPX4 ligand.

While targeting the catalytic U46 is a well-established strategy to abrogate GPX4 enzymatic activity, emerging evidence indicates that additional residues, such as C66,^25^ also contribute to regulating its function. Moreover, palmitoylation at C66 and C75 has been shown to enhance protein stability and dictate ferroptosis sensitivity,^40,41^ suggesting that solely targeting the catalytic center may not fully capture the pharmacological potential of GPX4 modulation. This complexity is further exemplified in patients with Sedaghatian-type spondylometaphyseal dysplasia (SSMD), who harbor the homozygous GPX4^R152H^ variant, a mutation associated with partial loss-of-function - likely due to altered PRMT5-mediated arginine methylation^42^ - while simultaneously rendering the protein more resistant to degradation.^43^ Beyond its primary peroxidase function, GPX4 acts as a moonlighting protein capable of forming enzymatically inactive structural assemblies with non-catalytic roles.^44,45^ Together, these findings underscore the importance of targeting both enzymatic and non-enzymatic functions of GPX4, thereby motivating the expansion of the GPX4 chemical toolbox from traditional inhibitors to degraders.^46,47^ However, existing GPX4 proteolysis-targeting chimeras (PROTACs) are largely derived from non-selective covalent inhibitors and often lack comprehensive proteome-wide validation of selectivity.^48-54^ Leveraging the exceptional selectivity profile of VK-3-189, we reasoned that its conversion into a PROTAC could provide a more precise tool for targeted GPX4 degradation.

To enable linker installation, the 2,6-dimethylmorpholine motif in VK-3-189 was replaced with a piperazine moiety as a handle for conjugation to a CRBN-recruiting ligand. This design ultimately led to the discovery of VK-7-91 as a potent GPX4-targeting PROTAC (Figure 5A). VK-7-91 retained potent cell-killing activity, with IC_50_ values of 12 nM in MDA-MB-231 cells (Figure S1A) and 16 nM in HT-1080 cells (Figure 5B). The latter was selected for subsequent degradation assays as it is a well-established and highly sensitive model for ferroptosis research. To evaluate the kinetics of GPX4 degradation, we performed a time-course experiment in HT-1080 cells treated with 0.5 μM VK-7-91. To prevent confounding cell death during the assay, cells were co-treated with 1.5 μM fer-1, a practice maintained for all subsequent degradation experiments unless otherwise specified. Degradation occurred gradually, with 54% reduction observed at 6 h and 75% achieved at 16 h (Figure 5C). The normalized degradation was quantified by integrating the intensities of both the upper and lower bands, with the former being consistent with the expected molecular weight of a covalent GPX4-PROTAC adduct. Further dose titration immunoblot analysis confirmed the potent degradation activity of VK-7-91. After 16 h of treatment, the compound achieved a maximal degradation (D_max_) of 87% at 333 nM, with a half-maximal degradation concentration (DC_50_) value of 1.3 nM (Figure 5D). Similar potency was observed at 6 h, yielding a D_max_ of 73% at 111 nM and a DC_50_ value of 4.0 nM (Figure 5E). We confirmed this process is both proteasome- and CRBN-dependent, as degradation was significantly rescued following pretreatment with the proteasome inhibitor MG132 or siRNA-mediated knockdown of CRBN (Figure S1B-D). Finally, to evaluate the activity of VK-7-91 across different cancer lineages, we tested the leukemia cell line MOLM-13, where GPX4 degradation remained evident (Figure S1E,F).

**Figure 5.**
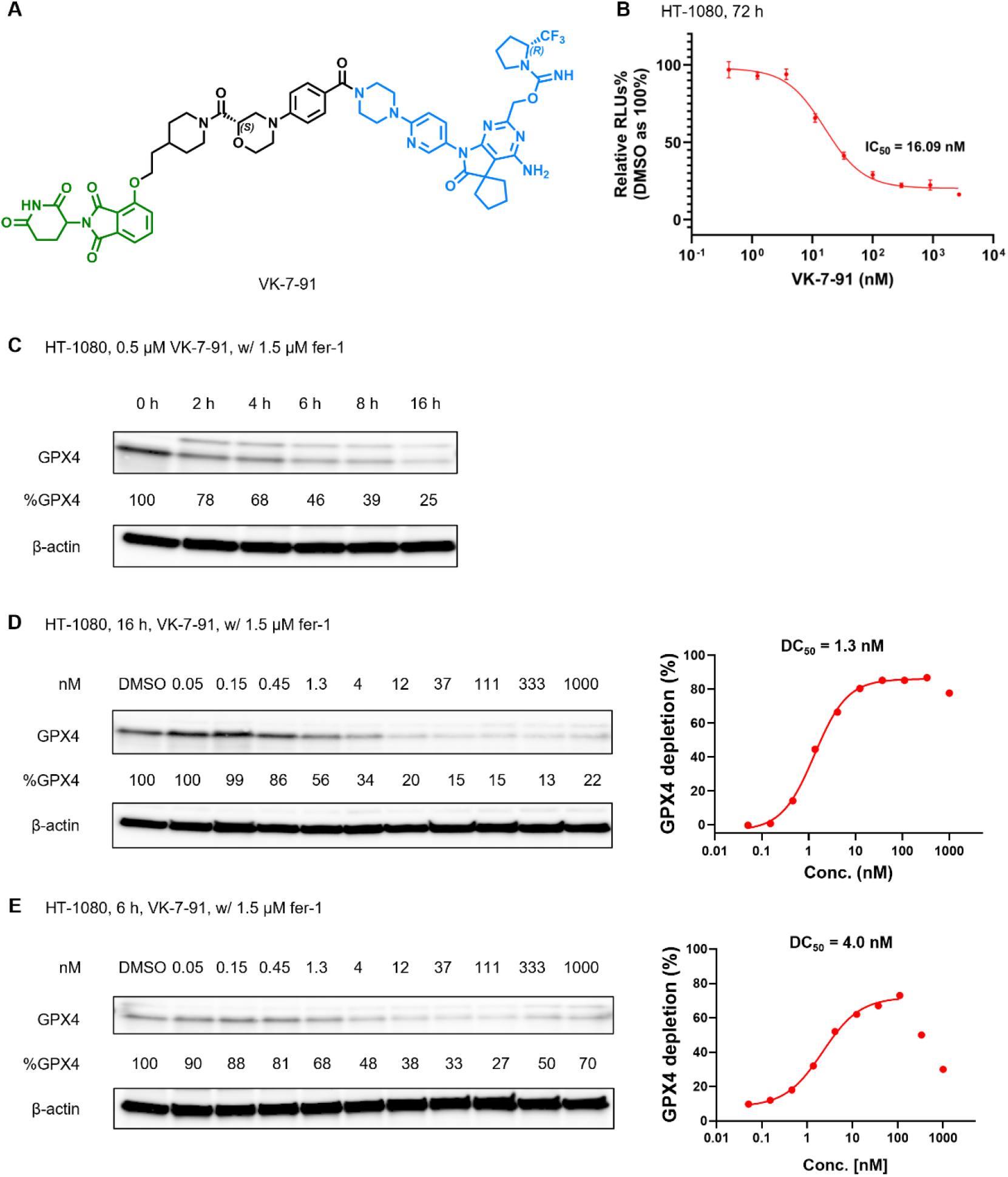
VK-7-91 is a potent GPX4 PROTAC. (A) Chemical structure of VK-7-91 highlighting the CRBN ligand (green), linker (black), and GPX4 ligand (blue). (B) Cell-killing effect following 72 h treatment with VK-7-91 in HT-1080 cells. Data are shown as mean ± s.d. (n = 3 biological replicates). (C) Immunoblot analysis of GPX4 levels in HT-1080 cells treated with 0.5 µM VK-7-91 over the indicated time points in the presence of 1.5 µM fer-1. The higher molecular weight band likely represents a covalent GPX4/VK-7-91 adduct. (D) Immunoblot analysis of GPX4 levels in HT-1080 cells treated with indicated concentrations of VK-7-91 in the presence of 1.5 µM fer-1 for 16 h or (E) 6 h.

Next, to rigorously validate the mechanism of action, we first synthesized the warhead-free analogue VK-7-90. As expected, this control was incapable of either inducing cell-killing effect or GPX4 reduction (Figure S2), confirming that the covalent warhead is indispensable for biological activity. We next sought to confirm CRBN-dependence using VK-8-92, which features an *N*-methylated glutarimide motif designed to abolish CRBN binding (Figure 6A). Paradoxically, VK-8-92 exhibited GPX4 degradation potency comparable to the parent PROTAC VK-7-91 (Figure 6B). We hypothesized that in this specific structural context, the methyl group transforms the intended CRBN ligand into a hydrophobic degron^55,56^ instead of merely serving as an inactive control. Supporting this, the introduction of a bulky benzyl group (VK-8-164) similarly maintained degradation activity, whereas the truncated, CRBN ligand-free analog VK-8-129 showed no effect (Figure S3). While the specific protein quality control-associated E3 ligase mediating this process remains to be identified, these findings underscore a novel covalent PROTAC mechanism^57^ where an ‘inactive’ ligand can bypass traditional E3 recruitment by functioning as a potent hydrophobic degron tag.

**Figure 6.**
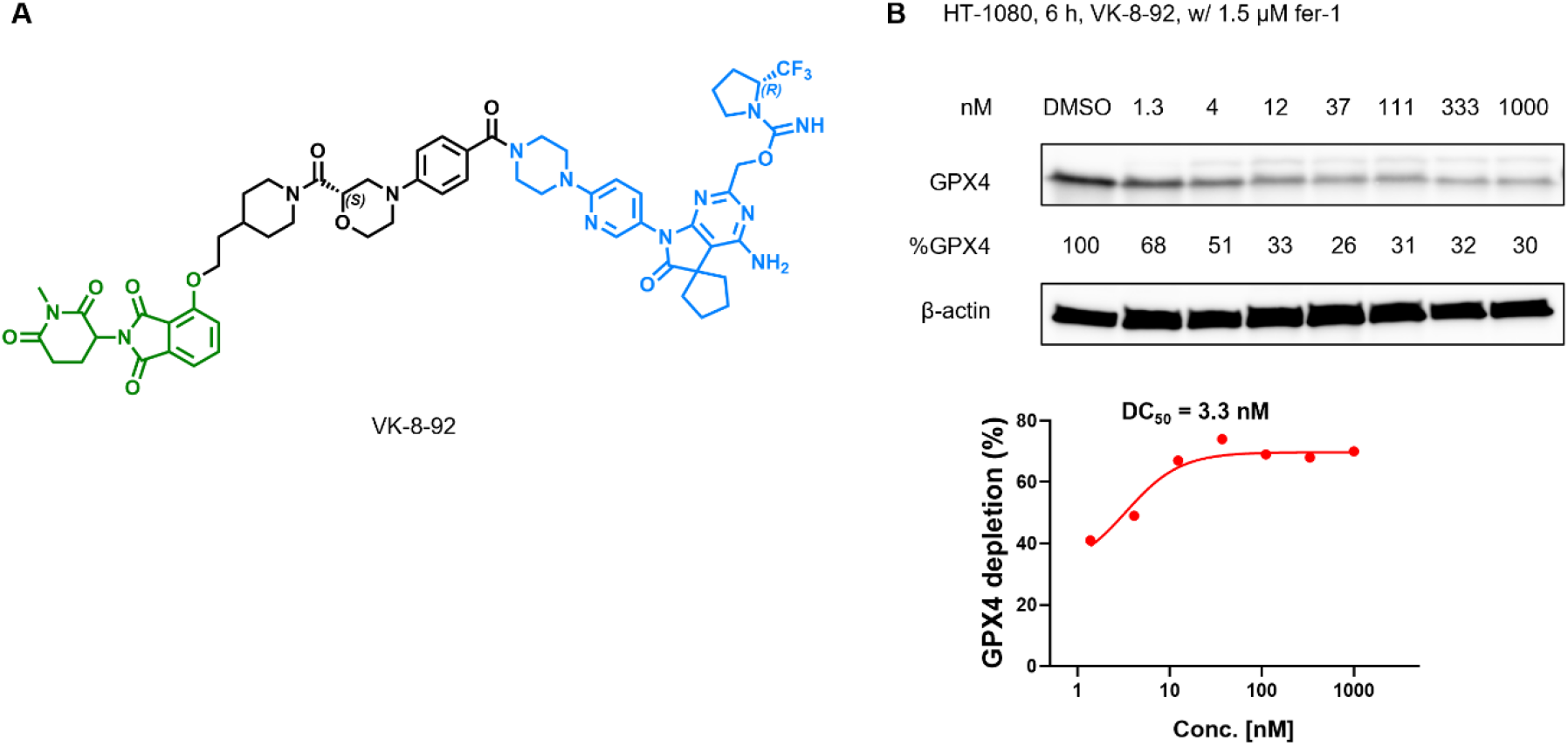
VK-8-92 induces GPX4 degradation independently of CRBN binding. (A) Chemical structure of VK-8-92. (B) Immunoblot analysis of GPX4 levels in HT-1080 cells treated with indicated concentrations of VK-8-92 for 6 h in the presence of 1.5 µM fer-1.

To assess the global degradation selectivity, we performed quantitative proteomic analysis of VK-7-91 and VK-8-92 in HT-1080 cells. As shown in Figure 7A and B (see also Data S1), following a 6 h treatment at 111 nM, both compounds exhibited exceptional proteome-wide selectivity for GPX4. Notably, only a single significant off-target, ODC1, was observed for VK-7-91, while VK-8-92 maintained a remarkably clean profile. Finally, we confirmed that this selective degradation translates into a functional ferroptosis-inducing phenotype. Consistent with the benchmark RSL3, treatment with either VK-3-189 or VK-7-91 (200 nM, 6 h) led to a robust accumulation of lipid hydroperoxides as quantified by flow cytometry. These effects were counteracted by co-treatment with fer-1 (Figure 8). Together, these results demonstrate that the inhibitor VK-3-189, along with its derived degraders VK-7-91 and VK-8-92, are highly precise chemical tools suitable for interrogating the biological roles and therapeutic potential of chemically induced GPX4 inhibition or depletion.

**Figure 7.**
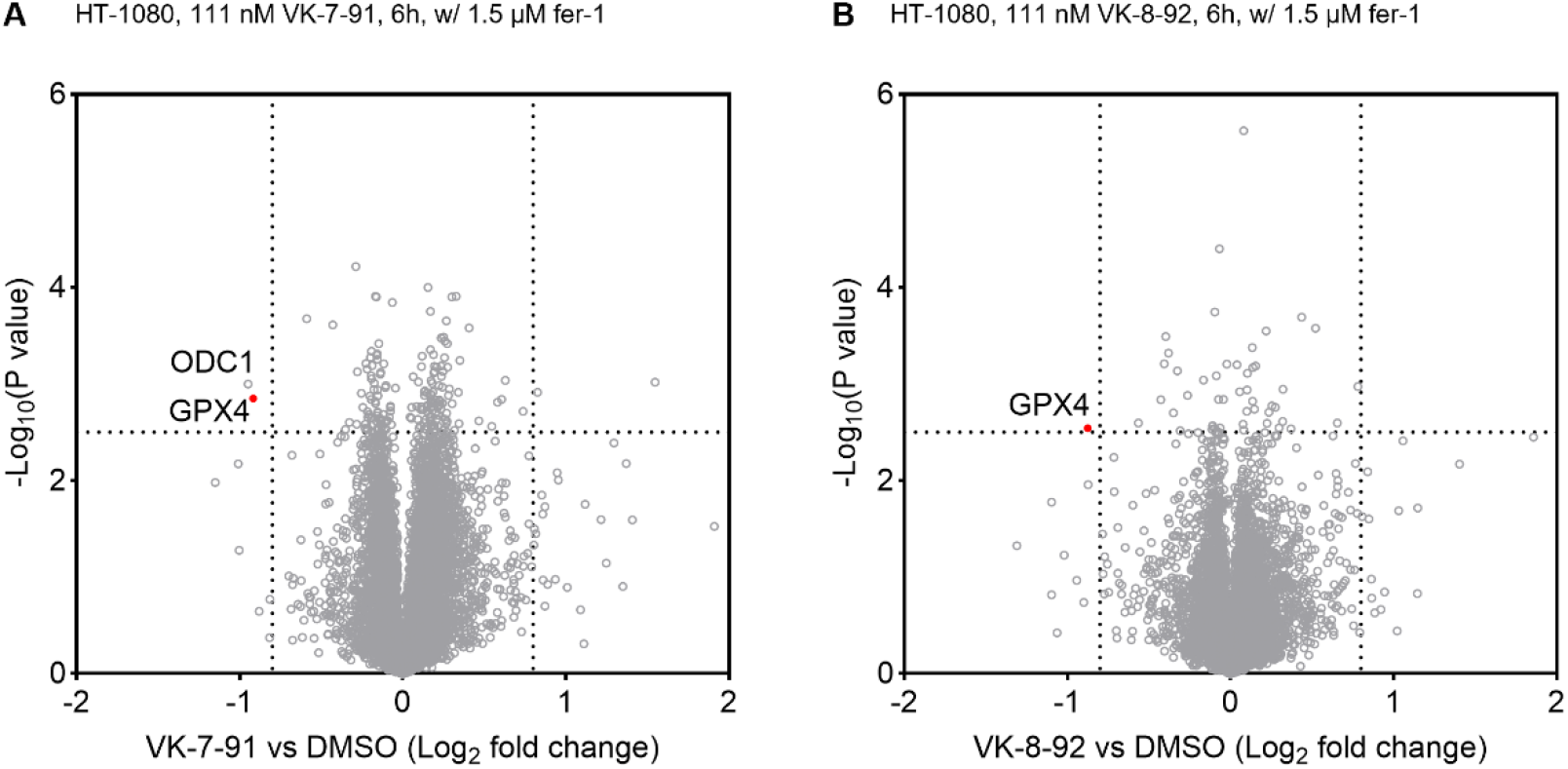
Proteome-wide degradation selectivity of VK-7-91 and VK-8-92. Global protein degradation profiles in HT-1080 cells treated for 6 h with 111 nM of either (A) VK-7-91 or (B) VK-8-92 in the presence of 1.5 µM Fer-1.

**Figure 8.**
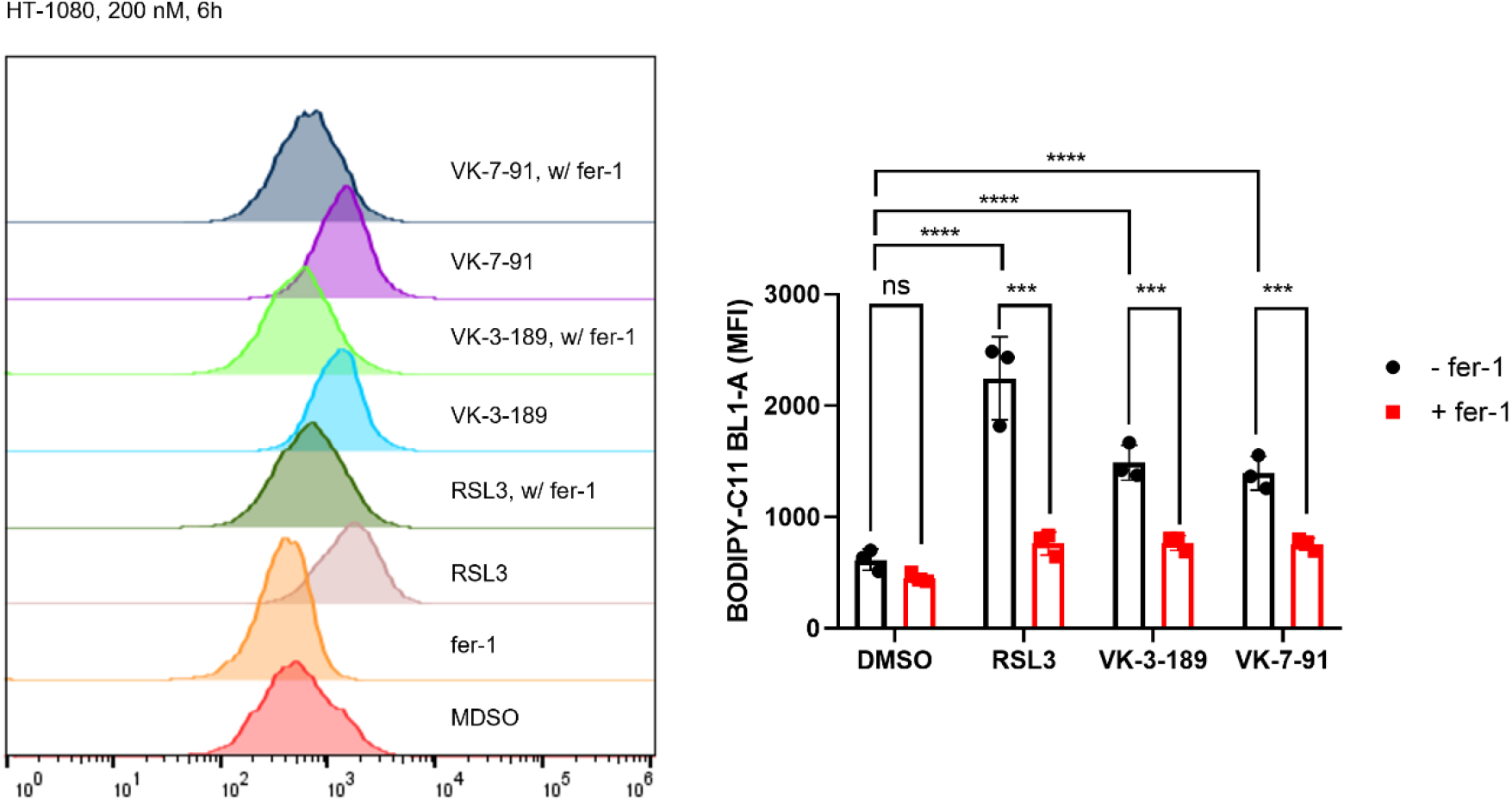
Assessment of lipid hydroperoxide accumulation. Treatment of HT-1080 cells with 200 nM RSL3, VK-3-189, or VK-7-91 (6 h) leads to the accumulation of lipid hydroperoxides, which is suppressed by co-treatment with 1.5 µM fer-1. Lipid hydroperoxide levels were quantified via flow cytometric analysis of C11-BODIPY 581/591 fluorescence. The x-axis shows the cell count. Data are plotted as the mean ± s.d. (n = 3 biological replicates). P values were determined by one-way ANOVA; **P < 0.01, ***P < 0.001, ****P < 0.0001. MFI, mean fluorescence intensity. ns, not significant.

## DISCUSSION

In this study, we leveraged a chemoproteomic approach to characterize a patented covalent GPX4 inhibitor featuring a novel pyrimidinylmethyl isourea warhead. From a chemical perspective, GPX4 remains a difficult-to-drug target due to its shallow active site, which necessitates a reliance on covalent targeting of the nucleophilic selenocysteine residue U46. Selenocysteine is generally considered more nucleophilic than cysteine because selenium is more polarizable than sulfur and selenocysteine possesses a significantly lower pKa (5.24) than cysteine (8.25), rendering the residue fully ionized at physiological pH.^58,59^ However, the residues surrounding U46 likely create a stringent environment that imposes specific requirements for warhead geometry, a structural constraint that may render the site incompatible with many common attenuated electrophiles.^14^ Consequently, successful targeting of GPX4 requires a finely tuned calibration between intrinsic warhead reactivity and precise spatial orientation to ensure proteome-wide selectivity - a challenge effectively addressed by the VK-3-189 molecule. This specific molecular architecture also highlights the pyrimidinylmethyl isourea motif as a ‘chemically programmable’ electrophile. By modulating substituents to tune steric and electronic effects, the leaving group ability can be precisely adjusted, suggesting broad utility for various other recalcitrant targets across the proteome.

Building upon the exceptional selectivity profile of VK-3-189, we further expanded the GPX4 chemical toolbox by transforming this inhibitor into a series of degraders. While several GPX4 PROTACs have been reported, they are largely derived from overly reactive covalent inhibitors and lack comprehensive, proteome-wide validation of their selectivity. In this work, we developed two degraders, VK-7-91 and VK-8-92, both of which exhibit outstanding selectivity in GPX4 degradation. Intriguingly, VK-8-92 - originally designed as a non-degrading negative control - retains degradation activity comparable to VK-7-91. We infer that its inactive CRBN ligand may be functioning as a hydrophobic degron, though further studies are warranted to confirm this mechanism. These tools are poised to address critical GPX4-related biological questions, particularly in the context of antitumor immunity, where emerging evidence suggests that GPX4 knockdown in immune cells may act as a double-edged sword.^4^

Nonetheless, this study has several limitations. First, in the cysteine-directed ABPP analysis, the GPX4 U46-containing peptide remained undetected, underscoring the need for improved methods to reliably capture selenoproteins. Second, the molecular interaction between VK-3-189 and GPX4 remains unclear. The relatively flat surface surrounding the catalytic selenocysteine in the active site,^18^ coupled with the bulky molecular size of VK-3-189 and its potent cellular activity, poses an apparent paradox. As discussed above, a membrane-associated state of GPX4 may provide a more defined binding environment for ligand engagement, warranting further structural investigation. Finally, during the preparation of VK-3-189-derived PROTACs, we occasionally observed that the installed warhead undergoes retro-addition in DMSO stock solutions, releasing a warhead-free pyrimidin-2-ylmethanol species and resulting in loss of GPX4 activity. This finding highlights the need for further chemical optimization to improve the stability of this warhead scaffold.

In conclusion, we characterized a new chemotype of covalent GPX4 inhibitor, VK-3-189, and developed two derived degraders, VK-7-91 and VK-8-92. Together, these molecules represent precise chemical tools for interrogating GPX4 biology and evaluating its therapeutic potential through both targeted inhibition and depletion.

## Author Contributions

^†^V.D.K., G.B., and C.M. contributed equally as co-first authors. *M.T., X.Z., and Z.T. are the corresponding authors. M.T. conceived the project and wrote the manuscript. M.T. supervised V.D.K.; M.T. and Z.T. supervised G.B.; X.Z. supervised C.M.; V.D.K. designed and synthesized the compounds; G.B. performed the majority of biological experiments and data analysis, with assistance from Z.X., M.L., and D.N.; C.M. conducted the chemoproteomic studies; H.G. performed CellTiter-Glo^®^ cell viability assays; Q.M. carried out the DTNB-based thiol reactivity experiments; J.W. and F.L. assisted with solubility and liver microsomal metabolic stability studies.

## Notes

The authors declare no competing financial interest.

## ACKNOWLEDGMENTS

The authors thank Dr. Yikai Wang and Dr. Jun Yu at Keen Therapeutics, Dr. Fu Gui and Dr. Li Jia at Brigham and Women’s Hospital, Dr. Scott Ficarro and Dr. Jarrod Marto at University of Virginia, Dr. Brian Fuglestad at Virginia Commonwealth University, and Dr. Martin Matzuk at Baylor College of Medicine, for their help. D.N. was supported by the NIH (R01CA255813) and the Henry and Emma Meyer Professorship in Molecular Genetics. Z.T. is a CPRIT Scholar in cancer research, and Z.T. thanks the CPRIT for research funding support (RR220039). M.T. is a CPRIT Scholar in cancer research and M.T. thanks the CPRIT for research funding support (RR220012).

## Abstract Graphic (For Tables of Contents Only)

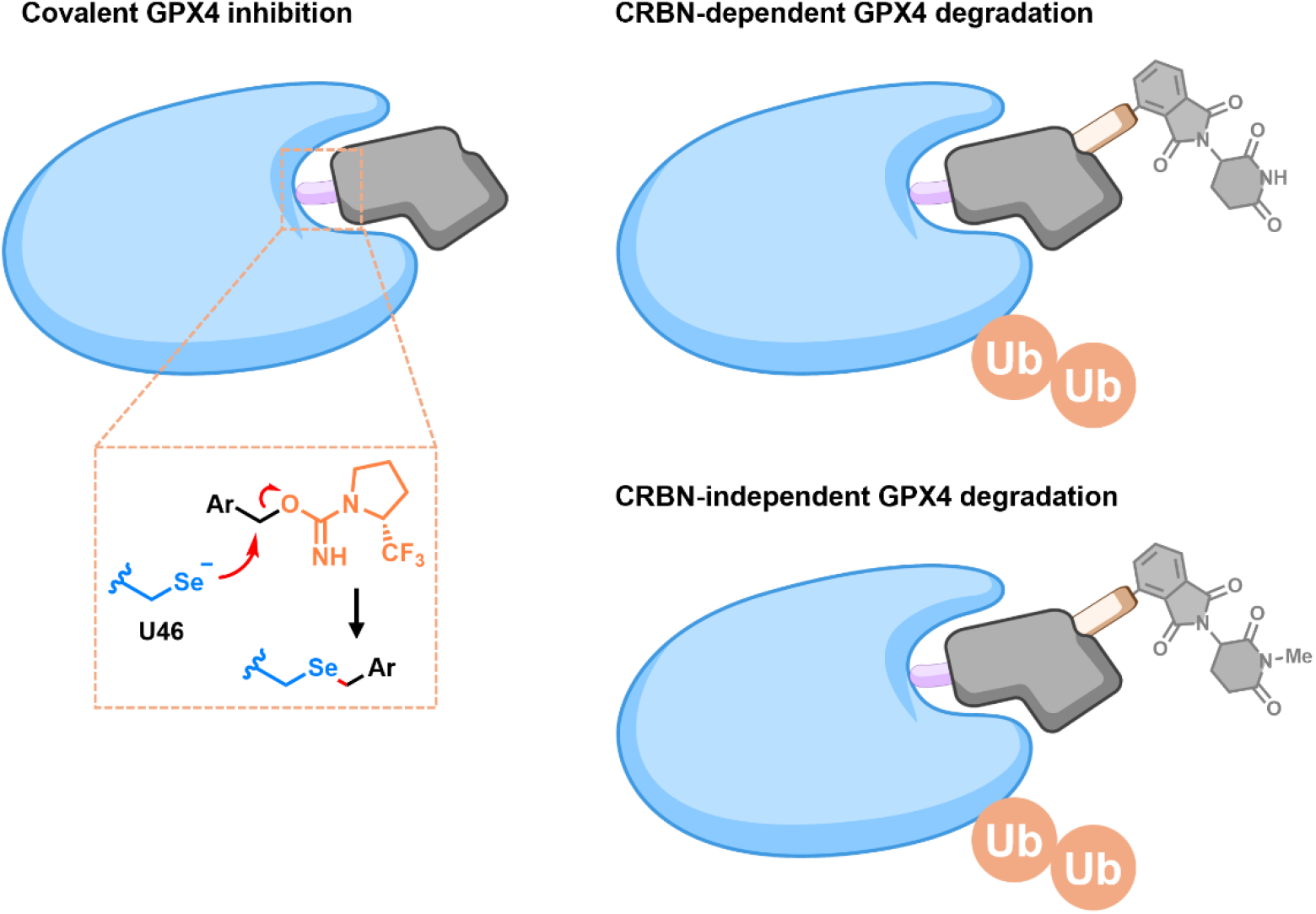

