## Supporting Information for "Chemoproteomic Characterization of GPX4 Covalent Ligands and Targeted Degradation"

**Table of Contents**

| | |
|---|---|
| Supporting Figures | Page 2 |
| Methods for Assays | Page 6 |
| Methods for Chemistry | Page 11 |
| NMR Spectra | Page 24 |

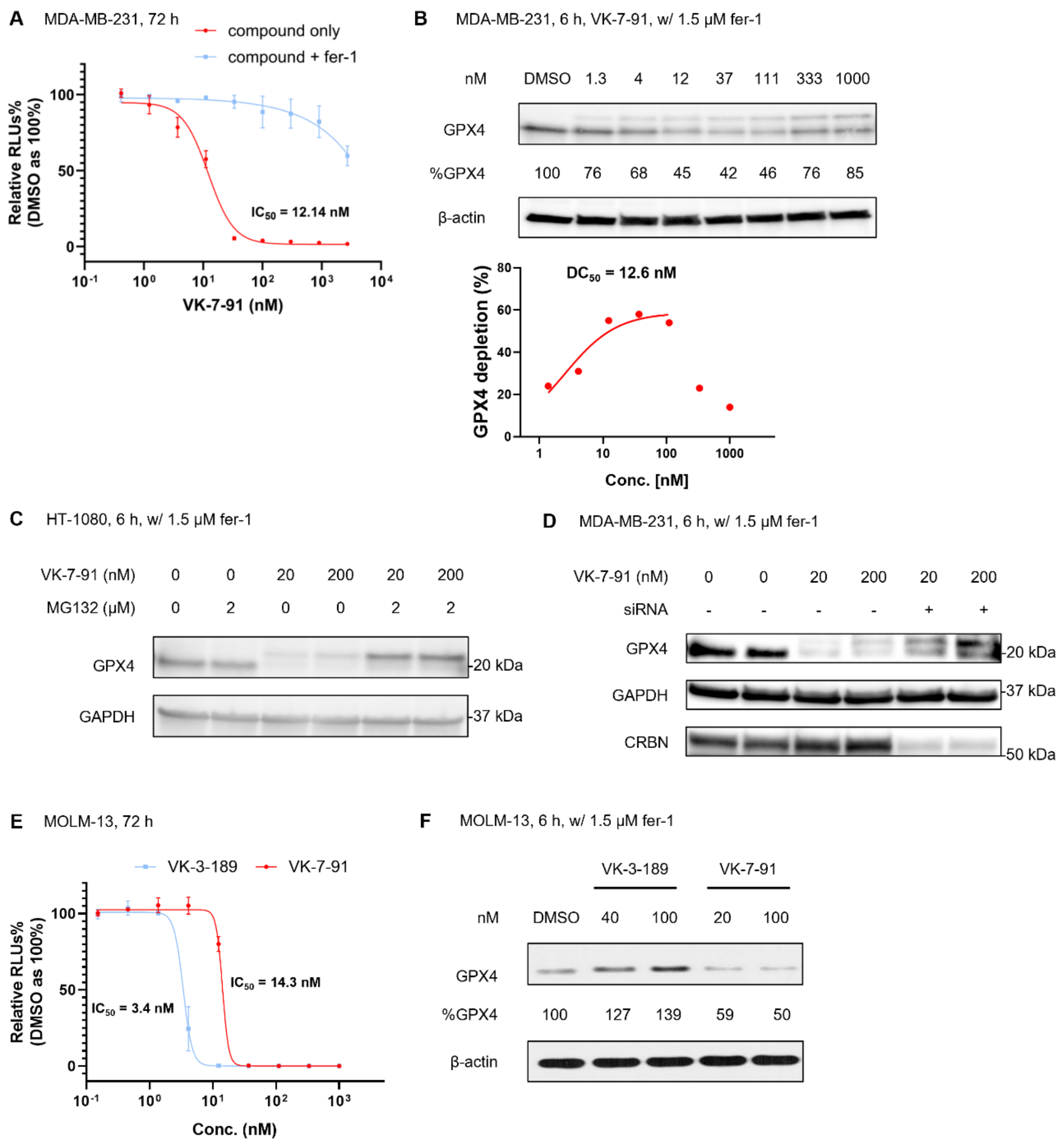


**Figure S1.** Validation of VK-7-91 as a potent and proteasome-dependent GPX4 degrader across multiple cancer cell lines. (A) Cell-killing effects following 72 h treatment with VK-7-91 in MDA-MB-231 cells, with rescue by 1.5 µM fer-1. Data are shown as mean ± s.d. (n = 3 biological replicates). (B) Immunoblot analysis of GPX4 levels in MDA-MB-231 cells treated with indicated concentrations of VK-7-91 for 6 h in the presence of 1.5 µM fer-1. (C) Immunoblot analysis in HT-1080 cells demonstrates that MG-132 (2 µM) prevents VK-7-91-induced GPX4 degradation, confirming a proteasome-dependent mechanism. (D) Immunoblot analysis in MDA-MB-231 cells demonstrates that siRNA-mediated knockdown of CRBN prevents VK-7-91-induced GPX4 degradation, confirming a CRBN-dependent mechanism. (E) Cell-killing effects following 72 h treatment with either VK-3-189 or VK-7-91 in MOLM-13 cells. Data are shown as mean ± s.d. (n = 3 biological replicates). (F) Immunoblot analysis of GPX4 levels in MOLM-13 cells treated with indicated concentrations of VK-7-91 or the control VK-3-189 for 6 h in the presence of 1.5 μM fer-1.

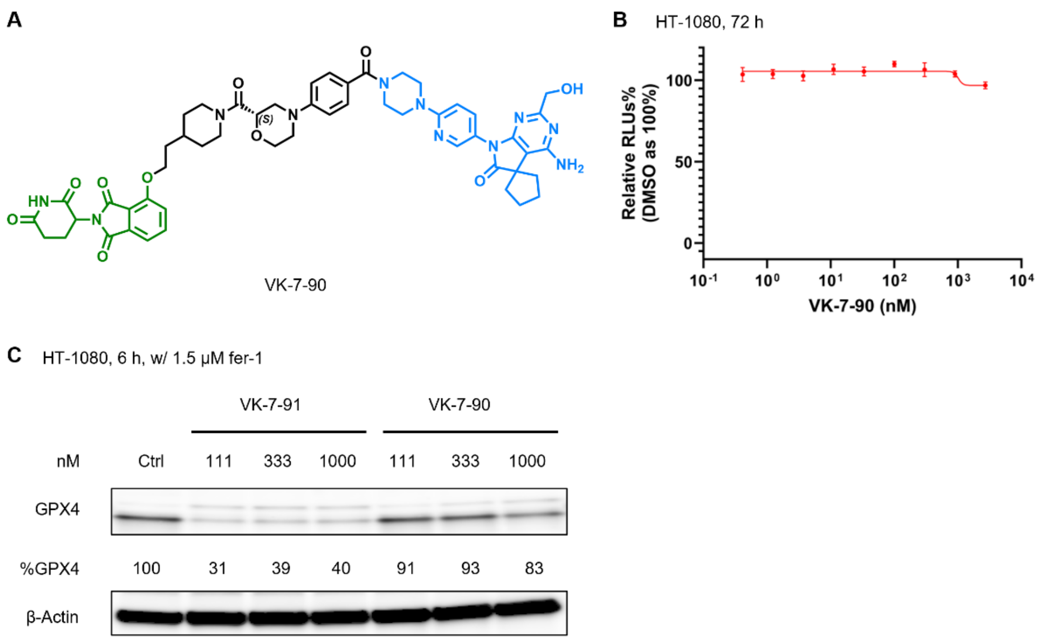


**Figure S2.** VK-7-90 serves as a non-degrading control for VK-7-91. (A) Chemical structure of VK-7-90. (B) Treatment of HT-1080 cells with VK-7-90 (72 h) shows no significant cell-killing effect, confirming its inactivity in engaging cellular GPX4. Data are shown as mean ± s.d. (n = 3 biological replicates). (C) Immunoblot analysis of GPX4 levels in HT-1080 cells treated with indicated concentrations of either VK-7-91 or VK-7-90 for 6 h in the presence of 1.5 μM fer-1, confirming the lack of degradation activity for VK-7-90. The higher molecular weight band observed with VK-7-90 likely corresponds to the long GPX4 variant, the precursor for mitochondrial import.

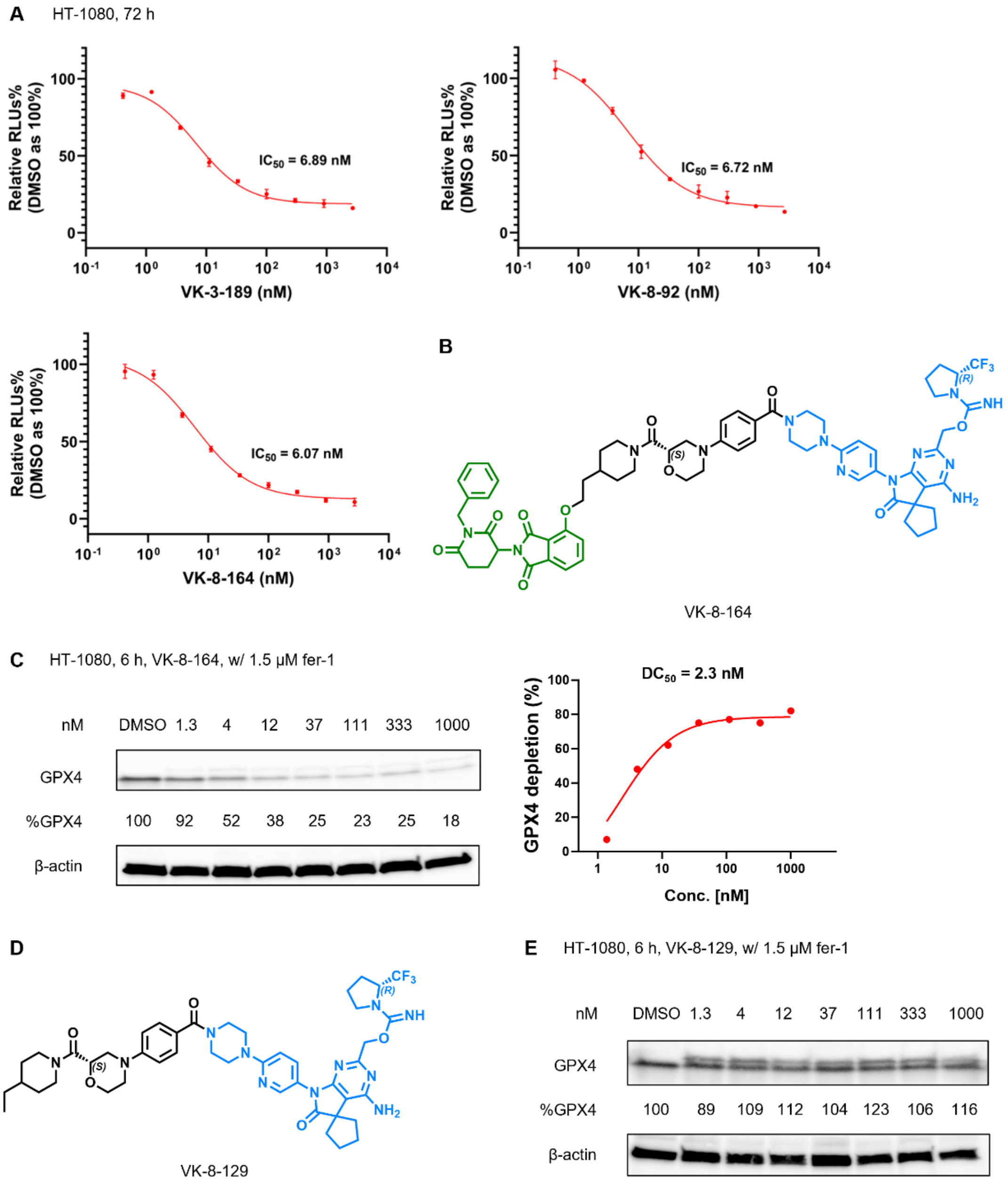


**Figure S3.** Functional evaluation of related GPX4 degraders independent of CRBN binding. (A) Cell-killing effect following 72 h treatment with VK-3-189, VK-8-92, or VK-8-164 in HT-1080 cells. Data are shown as mean ± s.d. (n = 3 biological replicates). (B) Chemical structure of VK-8-164. (C) Immunoblot analysis of GPX4 levels in HT-1080 cells treated with indicated concentrations of VK-8-164 for 6 h in the presence of 1.5 μM fer-1. (D) Chemical structure of VK-8-129. (E) Immunoblot analysis of GPX4 levels in HT-1080 cells treated with indicated concentrations of VK-8-129 for 6 h in the presence of 1.5 μM fer-1.

**Cell Culture**

The human breast adenocarcinoma cell line MDA-MB-231 (ATCC HTB-26) and the human fibrosarcoma cell line HT-1080 (ATCC CCL-121) were maintained in Dulbecco's Modified Eagle Medium (DMEM) (Gibco, cat# 11965092) supplemented with 10% fetal bovine serum (FBS) (Gibco, cat# A5256801) and 1% Penicillin-Streptomycin (Gibco, cat# 15140122). MOLM-13 cells were cultured under identical conditions, with the exception that RPMI-1640 medium (Gibco, cat# 11875093) was utilized as the basal medium. All cultures were incubated at 37 °C in a humidified atmosphere containing 5% $CO_2$. Cell lines were routinely screened for mycoplasma contamination using a PCR-based detection kit (Abcam, cat# AB289834) and remained negative throughout the study.

**Viability Assay**

MDA-MB-231 and HT-1080 cells were seeded into white 96-well plates (Costar, cat# 3917) at a density of 8,000 cells per well in 100 μL of complete medium. Following an 18 h initial incubation to allow for cell attachment, the medium was replaced with fresh medium containing test compounds at concentrations ranging from 0.41 nM to 2,700 nM (prepared as a 3-fold serial dilution). After 72 h of treatment, cell viability was quantified based on the ATP levels using the CellTiter-Glo® 2.0 Luminescent Assay (Promega, cat# G9242) according to the manufacturer's protocol. Luminescence signals were recorded using a BioTek Synergy Neo2 multi-mode plate reader. The half-maximal inhibitory concentration ($IC_{50}$) values were determined by fitting the normalized luminescence data to a four-parameter non-linear regression model (log[inhibitor] vs. response) using GraphPad Prism 11.0.

**Immunoblot Analysis**

Cells were harvested and lysed in RIPA buffer (Cell Signaling Technology, cat# 9806) supplemented with protease inhibitors (Sigma, cat# 11836153001). Protein concentrations were determined using a BCA protein assay kit (Thermo Fisher Scientific, cat# 23227). Equal amounts of protein were resolved by SDS-PAGE using 4-15% Criterion TGX Stain-Free Gels (Bio-Rad, cat# 5678084) and subsequently transferred to nitrocellulose (NC) membranes (Bio-Rad, cat# 1704271). The membranes were blocked and then incubated with primary antibodies overnight at 4 °C. The following primary antibodies were used: anti-GPX4 (Abcam, cat# ab41787, 1:1000), and anti-β-actin (Cell Signaling Technology, cat# 3700S, 1:1000). Following primary incubation, membranes were treated with HRP-linked anti-rabbit IgG (Cell Signaling Technology, cat# 7074S), or HRP-linked anti-mouse IgG (Cell Signaling Technology, cat# 7076S) secondary antibodies for 1 h at room temperature. Protein bands were visualized using the Bio-Rad ChemiDoc Imaging System. The half-maximal degradation concentration ($DC_{50}$) values were determined by fitting the normalized densitometry data to a four-parameter non-linear regression model (log[inhibitor] vs. response) using GraphPad Prism 11.0.

**CRBN Knockdown and Protein Degradation Rescue Assay**

MDA-MB-231 cells underwent reverse transfection with siRNA using Lipofectamine RNAiMAX (Invitrogen, cat# 3778075) according to the manufacturer's protocol. Briefly, siRNA lipid complexes were prepared directly in a 12-well plate by diluting a mixture of two CRBN-targeting siRNAs (hs.Ri.CRBN.13.3 and hs.Ri.CRBN.13.4; Integrated DNA Technologies; 20 pmol each) in 200 µL Opti-MEM reduced serum medium (Gibco, cat# 31985062). To this mixture, 2.2 µL Lipofectamine RNAiMAX was added, then the complexes were incubated for 20 min at room temperature to allow for assembly. Subsequently, 200,000 cells were seeded into each well in 800 µL of antibiotic-free DMEM (Gibco, cat# 11965092) supplemented with 10% FBS (Gibco, cat# A5256801). This resulted in a final transfection volume of 1 mL and a total siRNA concentration of 40 nM. The cells were gently mixed and incubated at 37 °C in a $CO_2$ atmosphere for 40 h prior to subsequent pharmacological treatment.

**Cysteine-Directed Activity-Based Protein Profiling (ABPP)**

Cells were plated on 10 cm plates to ~70% confluency and treated with the indicated compounds for 2 h at 37 ˚C. Following treatment, cells were harvested, washed with PBS, and lysed via probe sonification (40% intensity, 3 rounds, 5 pulses/round). Protein concentration was normalized to 1 mg/mL using a DC assay, and 500 µL of the normalized lysates were labeled with 100 µM desthiobiotin iodoactamide (DBIA) for 1 h at room temperature. Protein cleanup was facilitated using a adding a 1:1 mixture of hydrophobic:hydrophilic Sera-Mag SpeedBeads (100 µL). Lysates were incubated with the beads for 5 min (1,000 rpm) at room temperature, followed by protein precipitation through the addition of 1 mL absolute ethanol and further incubation. After magnetic separation using a DynaMag-2 magnet, the protein-bound beads were resuspended in 500 µL of 2 M urea/PBS, reduced with 25 µL of 10 mM dithiothreitol (DTT) at 65 °C for 15 min, and alkylated with 25 µL of iodoacetamide (400 mM in HPLC-grade water) at 37 °C for 30 min in the dark. The beads were then washed three times with 1 mL of absolute ethanol, resuspended in 200 µL of PBS, and digested with 2 µg of trypsin at 37 °C for 16 h. The resulting peptide supernatant was collected and enriched using 50 µL of streptavidin agarose in 300 µL of ABPP wash buffer (50 mM TEAB, 150 mM NaCl, 0.2% NP-40) with rotation at room temperature for 2 h. The beads were subsequently washed with ABPP wash buffer (3×1 mL), PBS (3×1 mL), and HPLC-grade water (3×1 mL) within a BioSpin column. Peptides were eluted off the beads with 300 µL of 50% acetonitrile/0.1% formic acid in HPLC-grade water, dried via SpeedVac, and resuspended in 100 µL of 100 mM TEAB/30% acetonitrile. For tandem mass tag (TMT) labeling, 3 µL of TMT tags were added to each sample for 1 h at room temperature, then quenched with 3 µL of a 5% hydroxylamine for 15 min. Samples were acidified with 5 µL of formic acid prior to pooling. Final desalting and fractionation were performed using a Pierce™ High pH Reversed-Phase Peptide Fractionation Kit (Thermo Fisher Scientific, Cat# 84868). Peptide samples were loaded onto a spin column and desalted by washing with HPLC-grade water containing 0.1% formic acid. This was followed by fractionation using fifteen increments of an increasing acetonitrile gradient in 10 mM $NH_4HCO_3$. Every 5th fraction was subsequently pooled and concentrated, resulting in five distinct fractions. Finally, peptides were analyzed by LC-MS/MS as described below.

**Competitive ABPP with Alkyne Enrichment**

Cells were plated on a 10 cm plate to ~70% confluency and treated with the indicated compounds for 2 h at 37 ˚C. Following treatment, cells were harvested, washed, and resuspended in PBS prior to lysis via probe sonification (40% intensity, 3 rounds, 5 pulses/round). Protein concentrations were normalized to 2 mg/mL by a DC assay. The cleared lysates were then tagged with a desthiobiotin-PEG3-azide probe (Sigma-Aldrich, cat# 902020) via a CuAAC reaction. A pre-mixed cocktail was added to reach a final volume of 500 µL with the final concentrations: 100 µM desthiobiotin-PEG3-azide, 100 µM tris(benzyltriazolylmethyl)-amine (TBTA), 1 mM $CuSO_4$, 1 mM freshly dissolved tris(2-carboxyethyl)phosphine (TCEP). The reaction was incubated for 1 h at room temperature with agitation. Protein samples were precipitated by adding 600 µL MeOH and 200 µL $CHCl_3$, followed by centrifugation (10 min, 4 ˚C, 21,300 *g*). The upper and lower solvent layers were discarded to isolate the protein disc. Proteins were washed by resuspension in 500 µL methanol via sonication (40% intensity, 1 round, 5 pulses/round) and collected by centrifugation (10 min, 4 ˚C, 21,300 *g*). The resulting pellet was air-dried at room temperature and then resuspended in 500 µL of freshly made 8 M urea in PBS supplemented with 10 µL of 10% SDS via sonication (40% intensity, 1 round, 5 pulses/round). For reduction, 25 µL of a 200 mM DTT solution in water was added and incubated at 65°C for 15 min, followed by alkylation with 25 µL of a 400 mM iodoacetamide solution in water at 37°C for 30 min in the dark. The samples were then supplemented with 100 µL of 10% SDS and transferred to a 15 mL tube containing 5 mL PBS. For enrichment, 50 µL of streptavidin beads per sample were washed, added to each sample, and rotated for 1.5 h at room temperature. The beads were subsequently washed with 0.2% SDS in PBS (2×1 mL), PBS (2×1 mL), HPLC water (2×1 mL), and 100 mM TEAB (1×1 mL). After aspiration of the supernatant, the beads were resuspended in 70 µL of 1M urea in 100 mM TEAB and digested with 2 µg of Trypsin/LysC at 37 °C for 16 h. The resulting peptide supernatant was collected and combined with 25 µL acetonitrile and 5 µL of TMT labels for 1 h at room temperature. The TMT labeling was quenched with 6 µL of 5% hydroxylamine for 15 min at room temperature and acidified with 5 µL of formic acid prior to pooling. The combined samples were dried via SpeedVac, desalted using a Sep-Pak C18 cartridge (Waters, cat# WAT054955), and the resulting elution was concentrated to dryness. Finally, peptides were analyzed by LC-MS/MS as described below.

**Global Proteomics**

Cells were plated on a 6-well plate to ~70% confluency and treated with the indicated compounds for 6 h at 37 ˚C. Following treatment, cells were harvested, washed with PBS, and lysed in 100 µL of PBS with complete protease inhibitor cocktail (Sigma-Aldrich, cat#: 11873580001) via sonication (40% intensity, 3 rounds, 5 pulses/round). Protein concentration was determined using a DC assay (BioRad, cat# 5000112). A total of 100 µg of protein in 100 µL of lysis buffer was denatured with 8 M urea. For reduction, 5 µL of 200 mM DTT stock solution in water was added, and the mixture was heated to 65 °C for 15 min, followed by alkylation with 5 µL of 400 mM iodoacetamide stock solution in water and incubating for 30 min at 37 °C in the dark. The samples were diluted with 300 µL of PBS and digested with 2 µg of Trypsin/LysC (Promega ,cat# V5071) at 37 °C for 16 h. For TMT labeling, aliquots containing approximately 8.5 µg peptide in 35 µL of solution was incubated with 9 µL of acetonitrile and 5 µL of TMT isobaric label reagent (Thermo Fisher Scientific, cat# 90110) for 1 h at room temperature. The reaction was quenched with 6 µL of 5% solution of hydroxylamine for 15 min at room temperature and acidified with 2.5 µL of formic acid prior to pooling. Desalting and fractionation were performed using a Pierce™ High pH Reversed-Phase Peptide Fractionation Kit (Thermo Fisher Scientific, cat# 84868). Briefly, the pooled peptide sample was loaded onto a spin column and desalted by washing with HPLC-grade water containing 0.1% formic acid, followed by fractionation using 30 increments of an increasing acetonitrile gradient in 10 mM $NH_4HCO_3$. Every 10th fraction was combined and concentrated, resulting in 10 distinct fractions for analysis. Peptides were analyzed by LC-MS using an Orbitrap Eclipse Tribid MS coupled with a Vanquish Neo UHPLC system. Peptides were loaded onto an EASY-Spray HPLC column (C18, 2 μm particle size, 75 μm inner diameter, and 250 mm length) and eluted at a flow rate of 0.25 μL/min using the following gradient: 5% buffer B (80% $CH_3CN$ with 0.1% FA) in buffer A (water with 0.1% FA) from 0-15 min, 5% to 45% buffer B from 15-155 min, and 45%-100% buffer B from 155-180 min. Voltage of the nano-LC electrospray ionization source was set to 1.5 kV. The analysis started with a MS1 master scan (Orbitrap analysis; resolution 60,000; m/z range 375-1600; RF lens 30%; standard automatic again control (AGC) target; auto maximum injection time). For MS2 analysis, initial precursor ions were isolated by the quadrupole with an isolation window of 0.7 and then subjected to higher-energy collisional dissociation (HCD) in the ion trap (stand AGC; collision energy 30%; maximum injection time 35 ms). After each MS2 spectrum, synchronous precursor selection (SPS) chose up to 10 MS2 fragment ions for MS3 analysis. These precursors were once again fragmented by HCD and analyzed by the Orbitrap (AGC 250%, collision energy 55%; maximum injection time 200 ms; resolution 60,000). The raw data was collected using Xcalibur (version 4.5.445.18).

**Thiol Reactivity Assay**

To assess thiol reactivity, 50 μM sample of DTNB was incubated with 200 μM tris(2-carboxyethyl)phosphine (TCEP) in 20 mM sodium phosphate buffer (pH 7.4) for 5 min at room temperature to generate $TNB^{2-}$. Test compounds (200 μM) were then added to the mixture, and absorbance at 412 nm was monitored immediately at 37 °C using a BioTek Synergy Neo2 multi-mode plate reader. Absorbance measurements were recorded every 15 min over a 7 h period. All assays were performed in triplicate in 96-well plates. To account for potential interference, the background absorbance of each compound was subtracted by measuring its absorbance under identical conditions in the absence of DTNB. The reaction kinetics were analyzed using a second-order rate equation. The rate constant (k) was determined as the slope of the linear regression of $\ln([A][B_0]/[B][A_0])$, where $[A_0]$ and $[B_0]$ represent the initial concentrations of the compound (200 μM) and $TNB^{2-}$ (100 μM), respectively, while [A] and [B] represent the remaining concentrations at time t as determined by the spectrometric data.

**Lipid Peroxidation Assay**

HT-1080 cells were seeded in 24-well plates at a density of 100,000 cells per well and incubated overnight. The cells were then treated for 4 h at 37 °C with RSL3 (MedChemExpress, cat# HY-100218A), VK-3-189, or VK-7-91 (all at 200 nM), in the presence or absence of 1.5 μM fer-1 (MedChemExpress, cat# HY-100579). Following treatment, the cells were washed three times with serum-free DMEM prior to fluorescent dye labeling. To detect lipid hydroperoxide levels, cells were incubated with the C11 BODIPY 581/591 probe (Invitrogen, cat# D3861) at a final concentration of 2 μM in serum-free medium for 20 min in the dark. Post staining incubation, the cells were washed three times with serum-free cell culture medium and harvested via trypsinization with 100 μl 0.25% Trypsin-EDTA (Gibco, cat# 25200072). The reaction was quenched with 0.5 mL of complete DMEM culture medium, and the cells were collected by centrifugation at 1,200 rpm for 3 min. The resulting pellet was resuspended in Opti-MEM Medium (Gibco, cat# 11058021), filtered through a 70-μm nylon mesh, and analyzed using an Attune NxT Flow Cytometer (Invitrogen). Data analysis was performed using FlowJo V10 software.

**Kinetic Solubility Assessment**

To 196 μL of 1× PBS, 4 μL compound stock (10 mM in DMSO) was added. The solutions were incubated at 25 °C for 1 h and centrifuged at 15,000 *g* for 10 min at room temperature. The resulting supernatants were transferred to sample vials, and 2 μL aliquots were injected for LC-MS/MS analysis. All samples were analyzed using a Thermo Vanquish UHPLC coupled to a Thermo TSQ Quantis MS equipped with a heated electrospray ionization source. Test compounds were retained on a Phenomenex Luna C18 column (1 mm × 50 mm, 1.6 μm, Torrance, CA) at 40 °C and eluted using a water-acetonitrile mobile phase system (both containing 0.1% formic acid, v/v) at a flow rate of 0.15 mL/min. The total LC-MS/MS run time was 5 min, with signal recorded from 2 to 4 min. The mass spectrometer operated in positive ESI mode with an ion spray voltage of 3500 V. High-purity nitrogen was used as sheath (35 arb) and auxiliary (7 arb) gases, while high-purity argon served as the collision gas. Compounds were monitored via selected reaction monitoring (SRM) using the transitions *m/z* 589 → 322 for VK-3-188 and *m/z* 589 → 407 for VK-3-189. To quantify the dissolved compounds, standard curves were established by injecting serial dilutions of stock solutions prepared in 50% methanol-water.

**Liver Microsomal Metabolic Stability Assessment**

The metabolic stability of test compounds was evaluated in pooled human and mouse liver microsomes (HLM and MLM, respectively; XenoTech, Lenexa, KS). Incubations were conducted in 1× PBS (pH 7.4) containing 2 μM test compound, 0.5 mg protein/mL liver microsomes, and 1.0 mM freshly prepared NADPH in a total volume of 400 μL. The reactions were performed in duplicate at 37 °C. At 0, 30, and 60 min, 100 μL aliquots were collected and quenched with 100 μL of ice-cold methanol containing agomelatine (0.1 μM) as an internal standard. Following quenching, samples were vortex-mixed and centrifuged at 15,000 *g* for 15 min at 4 °C. A 2 μL aliquot of the resulting supernatant was injected for LC-MS/MS analysis using the method described above. The percentage of compound remaining at each time point was calculated by normalizing the analyte peak area to the peak area at $t = 0$. Apparent half-lives ($t_{1/2}$) were estimated by linear regression of the natural logarithm of the remaining percentage versus incubation time.

#### Synthesis of compounds, characterization, and spectra

##### General information

Unless otherwise noted, reagents and solvents were obtained from commercial suppliers and were used without further purification. $^1H$ NMR spectra were recorded on Bruker Avance III HD 600 MHz spectrometer, and chemical shifts are reported in parts per million (ppm) with reference to solvent signals [DMSO-$d_6$ (2.50 ppm)]. Coupling constants (*J*) are reported in Hz. Spin multiplicities are described as s (singlet), brs (broad singlet), d (doublet), t (triplet), and m (multiplet). Mass spectra were obtained on a Waters Acquity UPLC. Flash column chromatography was carried out using Teledyne ISCO CombiFlash system equipped with either a silica or C-18 column (from 4g up to 100 g). Purities of assayed compounds were in all cases greater than 95%, as determined by UPLC.

##### Abbreviations used

$Bu_3SnCH_2OTBS$, tert-Butyl-dimethyl-(tributylstannylmethoxy)silane; $CH_3CN$, acetonitrile; $Cu(OAc)_2$, copper(II) acetate; DCM, dichloromethane; DIPEA, N,N-diisopropylethylamine; DMF, N,N-dimethylformamide; DMSO, dimethyl sulfoxide; DTBPF, 1,1′-bis(di-tert-butylphosphino)ferrocene; EtOAc, ethyl acetate; h, hours; HATU, 1-[bis(dimethylamino)methylene]-1H-1,2,3-triazolo[4,5-b]pyridinium 3-oxid hexafluorophosphate; HCl, hydrochloric acid; $K_2CO_3$, potassium carbonate; LDA, lithium diisopropylamide; MeOH, methanol; $Na_2SO_4$, sodium sulphate; $NaHCO_3$, sodium bicarbonate; $NH_4Cl$, ammonium chloride; NMP, N-Methyl-2-pyrrolidone; PyCIU, N,N,N′,N′-Bis(tetramethylene)chloroformamidinium hexafluorophosphate; rt, room temperature; $SOCl_2$, thionyl chloride; TFA, trifluoroacetic acid; THF, tetrahydrofuran; $ZnCl_2$, zinc chloride.

##### Experimental details for individual compound synthesis

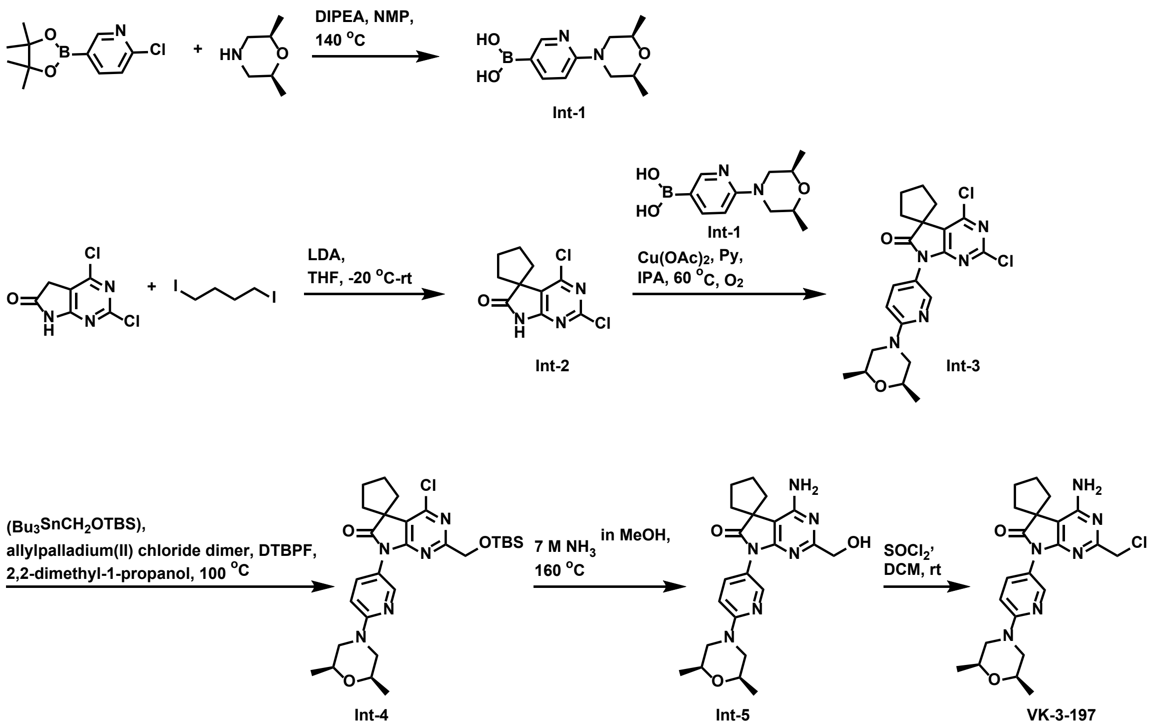


**(6-((2*S*,6*R*)-2,6-dimethylmorpholino)pyridin-3-yl)boronic acid (Int-1)**

To a solution of 2-chloro-5-(4,4,5,5-tetramethyl-1,3,2-dioxaborolan-2-yl)pyridine (5.0 g, 20.88 mmol) and (2*S*,6*R*)-2,6-dimethylmorpholine (2.6 g, 22.96 mmol) in NMP (30.0 mL) was added DIPEA (7.3 mL, 41.75 mmol). The resulting reaction mixture was stirred at 140 °C for 4 h. After completion of the reaction, the mixture was allowed to cool to rt, and 100 mL of ice-cold water was added. The resulting precipitate was collected by filtration, washed with cold water, and dried to afford **Int-1** (4.4 g, 89% yield) as a white solid. MS (ESI) for $C_{11}H_{17}BN_2O_3$ $[M+H]^+$: m/z calcd, 237.14; found, 237.16.

**2',4'-dichlorospiro[cyclopentane-1,5'-pyrrolo[2,3-d]pyrimidin]-6'(7'H)-one (Int-2)**

To a solution of 2,4-dichloro-5,7-dihydro-6*H*-pyrrolo[2,3-d]pyrimidin-6-one (3.0 g, 14.71 mmol) in dry THF (50.0 mL) was added LDA (16.2 mL, 32.35 mmol, 2 M in THF) dropwise at -20 °C under $N_2$ atmosphere. The resulting mixture was stirred at -20 °C for 1 h, followed by the addition of 1,4-diiodobutane (5.0 g, 16.18 mmol) at -20 °C. The reaction was allowed to warm to rt and stirred for 16 h. After completion of the reaction, the mixture was quenched with *sat. aq.* $NH_4Cl$ (15.0 mL) and diluted with water (100.0 mL). The aqueous layer was extracted with EtOAc (3 x 100.0 mL). The combined organic phases were dried over anhydrous $Na_2SO_4$, filtered, and concentrated under reduced pressure. The crude residue was purified via column chromatography (silica gel, eluted with 35% ethyl acetate in hexane) to give **Int-2** (2.24 g, 59% yield) as an off-white solid. MS (ESI) for $C_{10}H_9Cl_2N_3O$ $[M+H]^+$: m/z calcd, 258.02; found, 258.06.

**2',4'-dichloro-7'-(6-((2*S*,6*R*)-2,6-dimethylmorpholino)pyridin-3-yl)spiro[cyclopentane-1,5'-pyrrolo[2,3-d]pyrimidin]-6'(7'H)-one (Int-3)**

To a solution of **Int-2** (1.5 g, 5.81 mmol), **Int-1** (2.7 g, 11.62 mmol), and 4 Å MS (3.5 g) in isopropanol (20.0 mL) were added $Cu(OAc)_2$ (1.3 g, 6.97 mmol) and pyridine (1.4 mL, 17.43 mmol). The reaction mixture was stirred at 60 °C for 24 h under $O_2$ atmosphere. The reaction mixture was cooled to rt, filtered through a pad of Celite, and concentrated. The crude residue was purified via reverse-phase column chromatography on C-18 (eluted with 75% $CH_3CN$ in water) to afford **Int-3** (1.13 g, 43% yield) as a white solid. MS (ESI) for $C_{21}H_{23}Cl_2N_5O_2$ $[M+H]^+$: m/z calcd, 448.13; found, 448.14.

**2'-(((tert-butyldimethylsilyl)oxy)methyl)-4'-chloro-7'-(6-((2*S*,6*R*)-2,6-dimethylmorpholino)pyridin-3-yl)spiro[cyclopentane-1,5'-pyrrolo[2,3-d]pyrimidin]-6'(7'H)-one (Int-4)**

To a solution of **Int-3** (1.0 g, 2.23 mmol) and $Bu_3SnCH_2OTBS$ (1.4 g, 3.12 mmol) in 2,2-dimethyl-1-propanol (10.0 mL) were added allylpalladium(II) chloride dimer (40.0 mg, 0.11 mmol) and DTBPF (103.0 mg, 0.22 mmol). The reaction mixture was degassed by purging with $N_2$ for 15 minutes and stirred at 100 °C for 16 h. After completion of the reaction, the mixture was cooled to rt and filtered through a pad of Celite. The filtrate was concentrated under reduced pressure to provide a crude residue, which was purified via reverse-phase column chromatography on C-18 (eluted with 80-95% $CH_3CN$ in water) to afford **Int-4** (290.0 mg, 23% yield) as a light yellow solid. MS (ESI) for $C_{28}H_{40}ClN_5O_3Si$ $[M+H]^+$: m/z calcd, 558.27; found, 558.39.

**4'-amino-7'-(6-((2*S*,6*R*)-2,6-dimethylmorpholino)pyridin-3-yl)-2'-(hydroxymethyl)spiro[cyclopentane-1,5'-pyrrolo[2,3-d]pyrimidin]-6'(7'H)-one (Int-5)**

To a solution of **Int-4** (200.0 mg, 0.36 mmol) in MeOH (1.0 mL) was added ammonia (12.0 mL, 84.20 mmol, 7 M in MeOH). The reaction mixture was stirred at 160 °C for 24 h in hydrothermal autoclave reactor. The rection mixture was cooled to rt and the pressure was carefully released. The mixture was concentrated under reduced pressure, and the crude residue was purified via reverse-phase column chromatography on C-18 (eluted with 50% $CH_3CN$ in water) to afford **Int-5** (65.0 mg, 43% yield) as a white solid. $^1H$ NMR (600 MHz, DMSO-$d_6$) δ 8.12 (d, $J$ = 2.6 Hz, 1H), 7.57 (dd, $J$ = 9.1, 2.6 Hz, 1H), 6.93 (d, $J$ = 9.1 Hz, 1H), 6.54 (brs, 2H), 4.74 (t, $J$ = 5.9 Hz, 1H), 4.22 (d, $J$ = 5.9 Hz, 2H), 4.17 (dd, $J$ = 13.3, 2.3 Hz, 2H), 3.65-3.58 (m, 2H), 2.42 (dd, $J$ = 12.8, 10.6 Hz, 2H), 2.19-2.11 (m, 2H), 2.00-1.94 (m, 4H), 1.91-1.84 (m, 2H), 1.17 (d, $J$ = 6.2 Hz, 6H). MS (ESI) for $C_{22}H_{28}N_6O_3$ $[M+H]^+$: m/z calcd, 425.23; found, 425.29.

**4'-amino-2'-(chloromethyl)-7'-(6-((2*S*,6*R*)-2,6-dimethylmorpholino)pyridin-3-yl)spiro[cyclopentane-1,5'-pyrrolo[2,3-d]pyrimidin]-6'(7'H)-one (VK-3-197)**

To a solution of **Int-5** (5.0 mg, 11.78 µmol) in DCM (1.0 mL) was added $SOCl_2$ (3.0 µL, 41.22 µmol). The resulting mixture was stirred at rt for 6 h. The reaction mixture was concentrated under reduced pressure and purified via reverse-phase column chromatography on C-18 (eluted with 60% $CH_3CN$ in water) to afford **VK-3-197** (4.4 mg, 84% yield) as a white solid. $^1H$ NMR (600 MHz, DMSO-$d_6$) δ 8.11 (d, $J$ = 2.6 Hz, 1H), 7.57 (dd, $J$ = 9.1, 2.6 Hz, 1H), 6.94 (d, $J$ = 9.1 Hz, 1H), 6.71 (brs, 2H), 4.38 (s, 2H), 4.18 (dd, $J$ = 13.2, 2.4 Hz, 2H), 3.65-3.58 (m, 2H), 2.44 (dd, $J$ = 13.0, 10.6

Hz, 2H), 2.19-2.11 (m, 2H), 2.01-1.93 (m, 4H), 1.91-1.85 (m, 2H), 1.17 (d, *J* = 6.2 Hz, 6H). MS (ESI) for $C_{22}H_{27}ClN_6O_2$ $[M+H]^+$: m/z calcd, 443.20; found, 443.17.

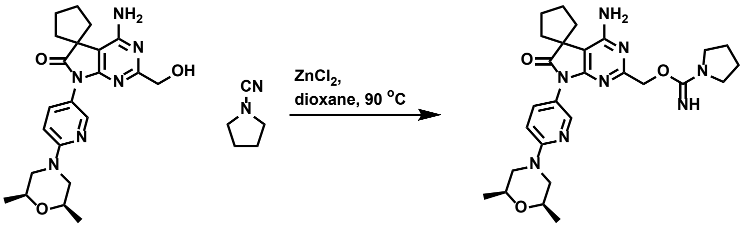


Int-5 VK-3-200

**(4'-amino-7'-(6-((2*S*,6*R*)-2,6-dimethylmorpholino)pyridin-3-yl)-6'-oxo-6',7'-dihydrospiro[cyclopentane-1,5'-pyrrolo[2,3-d]pyrimidin]-2'-yl)methyl pyrrolidine-1-carbimidate (VK-3-200)**

To a solution of **Int-5** (5.0 mg, 11.78 µmol) and pyrrolidine-1-carbonitrile (3.4 mg, 35.34 µmol) in dioxane (1.0 mL) was added $ZnCl_2$ (94.0 µL, 47.11 µmol, 0.5 M in THF) under $N_2$. The reaction mixture was stirred at 90 °C for 2 h. After completion of the reaction, the mixture was cooled to rt and concentrated under reduced pressure. The crude residue was purified via reverse-phase column chromatography on C-18 (eluted with 35% $CH_3CN$ in water) to afford **VK-3-200** (3.5 mg, 57% yield) as a white solid. $^1H$ NMR (600 MHz, DMSO-$d_6$) δ 8.58 (brs, 1H), 8.08 (d, *J* = 2.6 Hz, 1H), 7.56 (dd, *J* = 9.0, 2.6 Hz, 1H), 6.94 (d, *J* = 9.0 Hz, 1H), 6.74 (brs, 2H), 5.29 (s, 2H), 4.18 (dd, *J* = 13.1, 2.5 Hz, 2H), 3.66-3.56 (m, 2H), 3.32-3.28 (m, 4H), 2.43 (dd, *J* = 13.0, 10.6 Hz, 2H), 2.21-2.09 (m, 2H), 2.03-1.93 (m, 4H), 1.91-1.84 (m, 2H), 1.83-1.65 (m, 4H), 1.18 (d, *J* = 6.2 Hz, 6H). MS (ESI) for $C_{27}H_{36}N_8O_3$ $[M+H]^+$: m/z calcd, 521.30; found, 521.20.

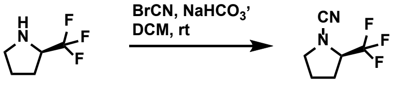


Int-6

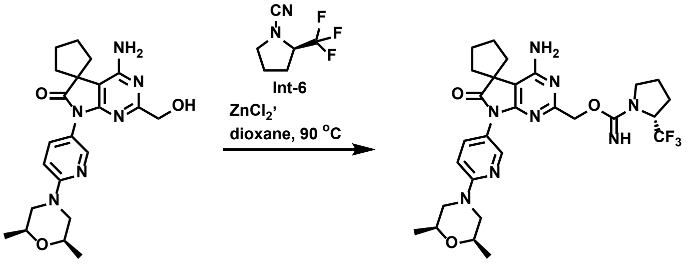


Int-5 VK-3-189

**(*R*)-2-(trifluoromethyl)pyrrolidine-1-carbonitrile (Int-6)**

To a solution of (*R*)-2-(trifluoromethyl)pyrrolidine (200.0 mg, 1.44 mmol) and $NaHCO_3$ (604.0 mg, 7.19 mmol) in DCM (2.0 mL) was added a solution of cyanogen bromide (304.5 mg, 2.88 mmol) in DCM (2.0 mL) at 0 °C. The reaction mixture was stirred at rt for 16 h. After completion of reaction, the mixture was quenched with water (10.0 mL) and extracted with DCM (2 x 20.0 ml). The combined organic layers were dried over anhydrous $Na_2SO_4$, filtered, and concentrated under reduced pressure. The crude residue was purified via column chromatography (silica gel, eluted with 30% ethyl acetate in hexane) to give **Int-6** (213.0 mg, 90% yield) as a colorless liquid. $^1$H NMR (600 MHz, DMSO-$d_6$) $\delta$ 4.59-4.50 (m, 1H), 3.58-3.53 (m, 1H), 3.46-3.40 (m, 1H), 2.23-2.15 (m, 1H), 1.99-1.91 (m, 2H), 1.88-1.80 (m, 1H).

**(4'-amino-7'-(6-((2*S*,6*R*)-2,6-dimethylmorpholino)pyridin-3-yl)-6'-oxo-6',7'-dihydrospiro[cyclopentane-1,5'-pyrrolo[2,3-d]pyrimidin]-2'-yl)methyl (*R*)-2-(trifluoromethyl)pyrrolidine-1-carbimidate (VK-3-189)**

Following the same procedure as compound **VK-3-200** by using **Int-6** instead of pyrrolidine-1-carbonitrile, compound **VK-3-189** (8.0 mg, 58% yield) was prepared as a white solid. $^1$H NMR (600 MHz, DMSO-$d_6$) δ 8.09 (d, $J$ = 2.6 Hz, 1H), 7.55 (dd, $J$ = 9.1, 2.6 Hz, 1H), 6.91 (d, $J$ = 9.1 Hz, 1H), 6.54 (brs, 2H), 5.76 (s, 1H), 5.03-4.77 (m, 2H), 4.58-4.43 (m, 1H), 4.21-4.12 (m, 2H), 3.64-3.56 (m, 2H), 3.30-3.18 (m, 2H), 2.42 (dd, $J$ = 13.0, 10.3 Hz, 2H), 2.18-2.09 (m, 2H), 2.01-1.93 (m, 4H), 1.90-1.72 (m, 6H), 1.17 (d, $J$ = 6.2 Hz, 6H). MS (ESI) for $C_{28}H_{35}F_3N_8O_3$ $[M+H]^+$: m/z calcd, 589.29; found, 589.32.

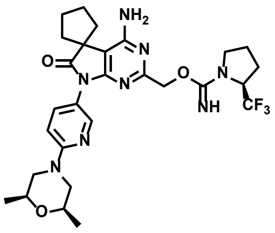

VK-3-188

**(4'-amino-7'-(6-((2*S*,6*R*)-2,6-dimethylmorpholino)pyridin-3-yl)-6'-oxo-6',7'-dihydrospiro[cyclopentane-1,5'-pyrrolo[2,3-d]pyrimidin]-2'-yl)methyl (*S*)-2-(trifluoromethyl)pyrrolidine-1-carbimidate (VK-3-188)**

Following the same procedure as compound **VK-3-200** by using (*S*)-2-(trifluoromethyl)pyrrolidine-1-carbonitrile instead of pyrrolidine-1-carbonitrile, compound **VK-3-188** (8.8 mg, 63% yield) was prepared as a white solid. $^1$H NMR (600 MHz, DMSO-$d_6$) δ 8.09 (d, $J$ = 2.6 Hz, 1H), 7.55 (dd, $J$ = 9.1, 2.6 Hz, 1H), 6.91 (d, $J$ = 9.1 Hz, 1H), 6.54 (brs, 2H), 5.76 (s, 1H), 5.03-4.77 (m, 2H), 4.58-4.43 (m, 1H), 4.21-4.12 (m, 2H), 3.64-3.56 (m, 2H), 3.30-3.18 (m, 2H), 2.42 (dd, $J$ = 13.0, 10.3 Hz, 2H), 2.18-2.09 (m, 2H), 2.01-1.93 (m, 4H), 1.90-1.72 (m, 6H), 1.17 (d, $J$ = 6.2 Hz, 6H). MS (ESI) for $C_{28}H_{35}F_3N_8O_3$ $[M+H]^+$: m/z calcd, 589.29; found, 589.41.

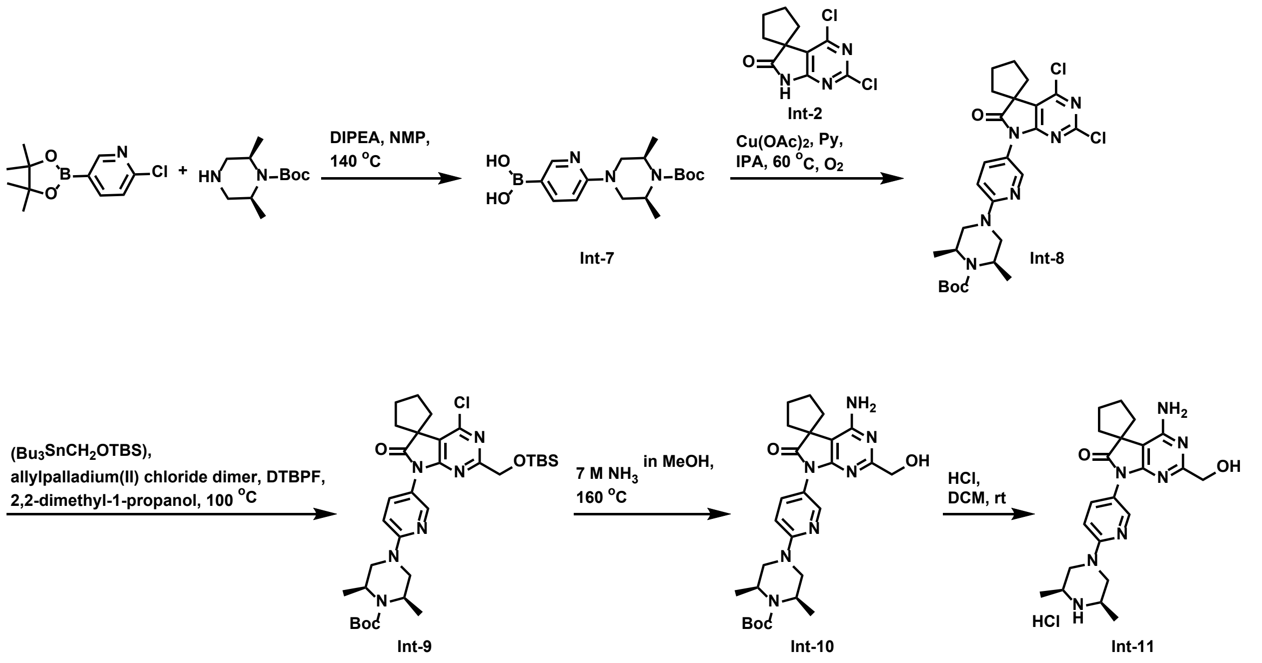


**(6-((3*S*,5*R*)-4-(tert-butoxycarbonyl)-3,5-dimethylpiperazin-1-yl)pyridin-3-yl)boronic acid (Int-7)**

Following the same procedure as compound **Int-1** by using tert-butyl (2*S*,6*R*)-2,6-dimethylpiperazine-1-carboxylate instead of (2*S*,6*R*)-2,6-dimethylmorpholine, compound **Int-7** (4.0 g, 71% yield) was prepared as a white solid. MS (ESI) for $C_{16}H_{26}BN_3O_4$ $[M+H]^+$: m/z calcd, 336.21; found, 336.21.

**tert-butyl (2*S*,6*R*)-4-(5-(2',4'-dichloro-6'-oxospiro[cyclopentane-1,5'-pyrrolo[2,3-d]pyrimidin]-7'(6'H)-yl)pyridin-2-yl)-2,6-dimethylpiperazine-1-carboxylate (Int-8)**

Following the same procedure as compound **Int-3** by using **Int-7** instead of **Int-1**, compound **Int-8** (600.0 mg, 32% yield) was prepared as a light brown solid. MS (ESI) for $C_{26}H_{32}Cl_2N_6O_3$ $[M+H]^+$: m/z calcd, 547.20; found, 547.30.

**tert-butyl (2*S*,6*R*)-4-(5-(2'-(((tert-butyldimethylsilyl)oxy)methyl)-4'-chloro-6'-oxospiro[cyclopentane-1,5'-pyrrolo[2,3-d]pyrimidin]-7'(6'H)-yl)pyridin-2-yl)-2,6-dimethylpiperazine-1-carboxylate (Int-9)**

Following the same procedure as compound **Int-4** by using **Int-8** instead of **Int-3**, compound **Int-9** (200.0 mg, 35% yield) was prepared as a light brown viscous oil. MS (ESI) for $C_{33}H_{49}ClN_6O_4Si$ $[M+H]^+$: m/z calcd, 657.34; found, 657.42.

**tert-butyl (2*S*,6*R*)-4-(5-(4'-amino-2'-(hydroxymethyl)-6'-oxospiro[cyclopentane-1,5'-pyrrolo[2,3-d]pyrimidin]-7'(6'H)-yl)pyridin-2-yl)-2,6-dimethylpiperazine-1-carboxylate (Int-10)**

Following the same procedure as compound **Int-5** by using **Int-9** instead of **Int-4**, compound **Int-10** (35.0 mg, 57% yield) was prepared as an off-white solid. MS (ESI) for $C_{27}H_{37}N_7O_4$ $[M+H]^+$: m/z calcd, 524.30; found, 524.39.

**4'-amino-7'-(6-((3*S*,5*R*)-3,5-dimethylpiperazin-1-yl)pyridin-3-yl)-2'-(hydroxymethyl)spiro[cyclopentane-1,5'-pyrrolo[2,3-d]pyrimidin]-6'(7'H)-one hydrochloride (Int-11)**

To a solution of **Int-10** (30.0 mg, 57.29 µmol) in DCM (1.0 mL) was added HCl (143.0 µL, 0.57 mmol, 4 M in dioxane) dropwise at 0 °C. The reaction mixture was stirred at rt for 6 h. The reaction mixture was concentrated under reduced pressure to give **Int-11** (23.0 mg, 87%) as an off-white solid. MS (ESI) for $C_{22}H_{29}N_7O_2$ $[M+H]^+$: m/z calcd, 424.25; found, 424.23.

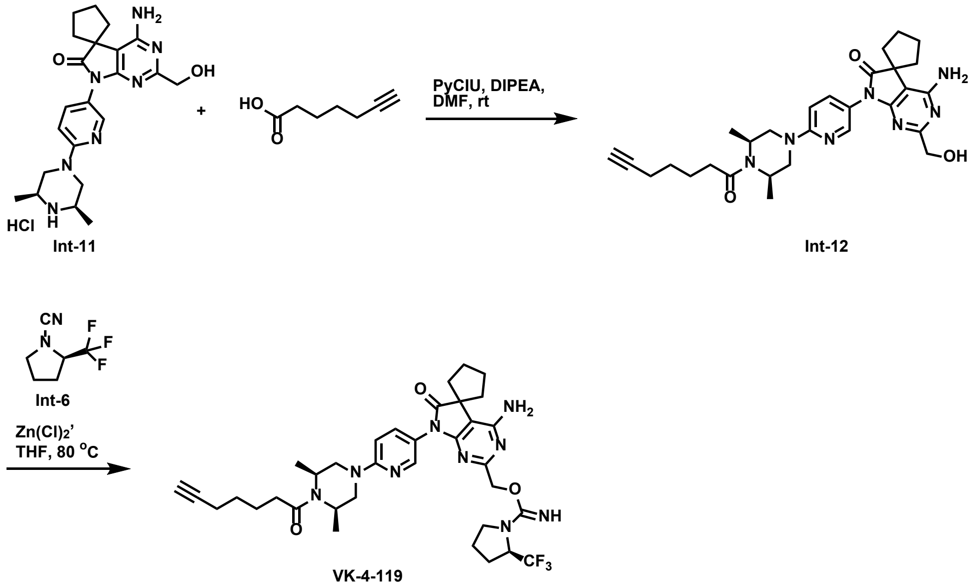


**4'-amino-7'-(6-((3*S*,5*R*)-4-(hept-6-ynoyl)-3,5-dimethylpiperazin-1-yl)pyridin-3-yl)-2'-(hydroxymethyl)spiro[cyclopentane-1,5'-pyrrolo[2,3-d]pyrimidin]-6'(7'H)-one (Int-12)**

To a solution of **Int-11** (10.0 mg, 21.74 µmol) and 6-heptynoic acid (2.7 mg, 21.74 µmol) in DMF (1.0 mL) were added DIPEA (19.0 µL, 0.11 mmol) and PyCIU (8.6 mg, 26.08 µmol). The reaction mixture was stirred at rt for 48 h. The reaction mixture was purified directly via reverse-phase column chromatography on C-18 (eluted with 50% $CH_3CN$ in water) to afford **Int-12** (5.0 mg, 57% yield) as a white solid. MS (ESI) for $C_{29}H_{37}N_7O_3$ $[M+H]^+$: m/z calcd, 532.30; found, 532.26.

**(4'-amino-7'-(6-((3*S*,5*R*)-4-(hept-6-ynoyl)-3,5-dimethylpiperazin-1-yl)pyridin-3-yl)-6'-oxo-6',7'-dihydrospiro[cyclopentane-1,5'-pyrrolo[2,3-d]pyrimidin]-2'-yl)methyl (*R*)-2-(trifluoromethyl)pyrrolidine-1-carbimidate (VK-4-119)**

To a solution of **Int-12** (5.0 mg, 9.40 µmol) and **Int-6** (15.4 mg, 94.04 µmol) in THF (1.0 mL) was added $ZnCl_2$ (188.0 µL, 94.04 µmol, 0.5 M in THF) under $N_2$. The reaction mixture was stirred at 80 °C for 12 h. After completion of the reaction, the mixture was cooled to rt and concentrated under reduced pressure. The crude residue was purified via reverse-phase column chromatography on C-18 (eluted with 70% $CH_3CN$ in water) to afford **VK-4-119** (5.0 mg, 85% yield) as a white solid. $^1$H NMR (600 MHz, DMSO-$d_6$) δ 8.08 (d, $J$ = 2.6 Hz, 1H), 7.55 (dd, $J$ = 9.1, 2.6 Hz, 1H), 6.98 (d, $J$ = 9.1 Hz, 1H), 6.53 (brs, 2H), 5.76 (s, 1H), 5.03-4.80 (m, 2H), 4.63-4.45 (m, 2H), 4.30-4.13 (m, 3H), 3.29-3.23 (m, 2H), 3.03-2.91 (m, 2H), 2.75 (t, $J$ = 2.6 Hz, 1H), 2.31-2.20 (m, 1H), 2.20-2.16 (m, 2H), 2.16-2.10 (m, 2H), 2.00-1.93 (m, 4H), 1.90-1.74 (m, 6H), 1.65-1.58 (m, 2H), 1.52-1.45 (m, 2H), 1.29-1.12 (m, 6H). MS (ESI) for $C_{35}H_{44}F_3N_9O_3$ $[M+H]^+$: m/z calcd, 696.36; found, 696.34.

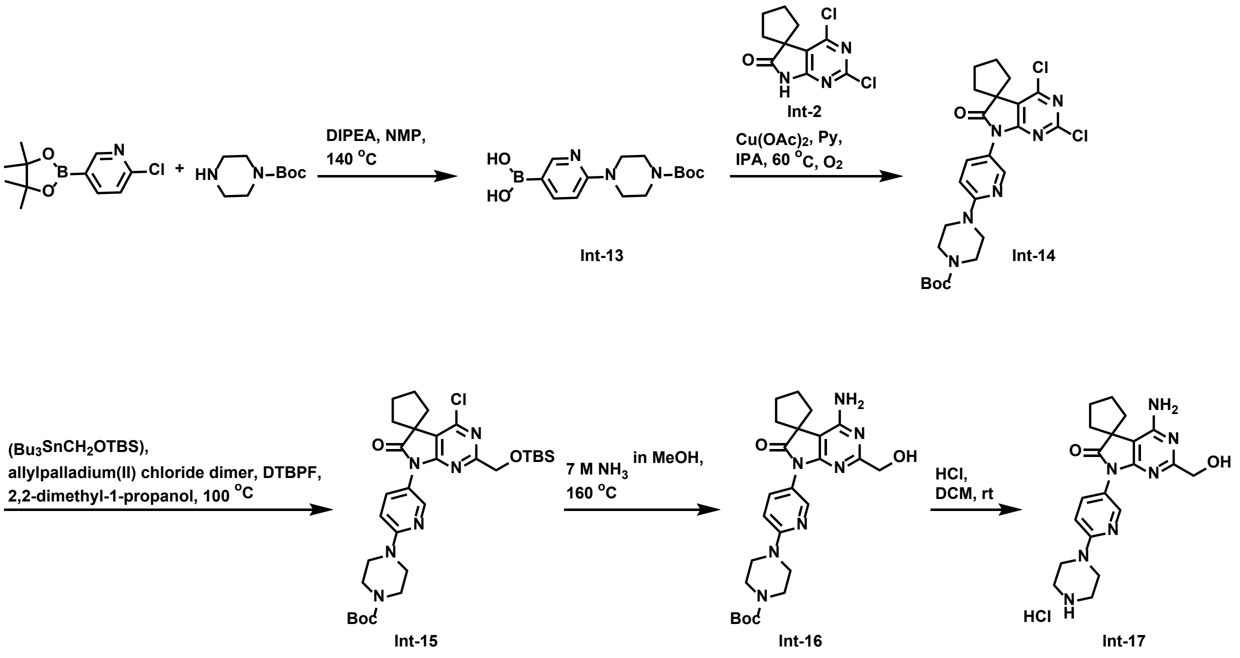


**(6-(4-(tert-butoxycarbonyl)piperazin-1-yl)pyridin-3-yl)boronic acid (Int-13)**

Following the same procedure as compound **Int-1** by using 1-boc-piperazine instead of (2*S*,6*R*)-2,6-dimethylmorpholine, compound **Int-13** (3.8 g, 98% yield) was prepared as a white solid. MS (ESI) for $C_{14}H_{22}BN_3O_4$ $[M+H]^+$: m/z calcd, 308.18; found, 308.25.

**tert-butyl 4-(5-(2',4'-dichloro-6'-oxospiro[cyclopentane-1,5'-pyrrolo[2,3-d]pyrimidin]-7'(6'H)-yl)pyridin-2-yl)piperazine-1-carboxylate (Int-14)**

Following the same procedure as compound **Int-3** by using **Int-13** instead of **Int-1**, compound **Int-14** (1.8 g, 29% yield) was prepared as a light brown solid. MS (ESI) for $C_{24}H_{28}Cl_2N_6O_3$ $[M+H]^+$: m/z calcd, 519.17; found, 519.26.

**tert-butyl 4-(5-(2'-(((tert-butyldimethylsilyl)oxy)methyl)-4'-chloro-6'-oxospiro[cyclopentane-1,5'-pyrrolo[2,3-d]pyrimidin]-7'(6'H)-yl)pyridin-2-yl)piperazine-1-carboxylate (Int-15)**

Following the same procedure as compound **Int-4** by using **Int-14** instead of **Int-3**, compound **Int-15** (600.0 mg, 31% yield) was prepared as an off-white solid. MS (ESI) for $C_{31}H_{45}ClN_6O_4Si$ $[M+H]^+$: m/z calcd, 629.30; found, 629.39.

**tert-butyl 4-(5-(4'-amino-2'-(hydroxymethyl)-6'-oxospiro[cyclopentane-1,5'-pyrrolo[2,3-d]pyrimidin]-7'(6'H)-yl)pyridin-2-yl)piperazine-1-carboxylate (Int-16)**

Following the same procedure as compound **Int-5** by using **Int-15** instead of **Int-4**, compound **Int-16** (300.0 mg, 67% yield) was prepared as an off-white solid. MS (ESI) for $C_{25}H_{33}N_7O_4$ $[M+H]^+$: m/z calcd, 496.27; found, 496.34.

**4'-amino-2'-(hydroxymethyl)-7'-(6-(piperazin-1-yl)pyridin-3-yl)spiro[cyclopentane-1,5'-pyrrolo[2,3-d]pyrimidin]-6'(7'H)-one hydrochloride (Int-17)**

Following the same procedure as compound **Int-11** by using **Int-16** instead of **Int-10**, compound **Int-17** (245.0 mg, 97% yield) was prepared as an off-white solid. MS (ESI) for $C_{20}H_{25}N_7O_2$ $[M+H]^+$: m/z calcd, 396.21; found, 396.28.

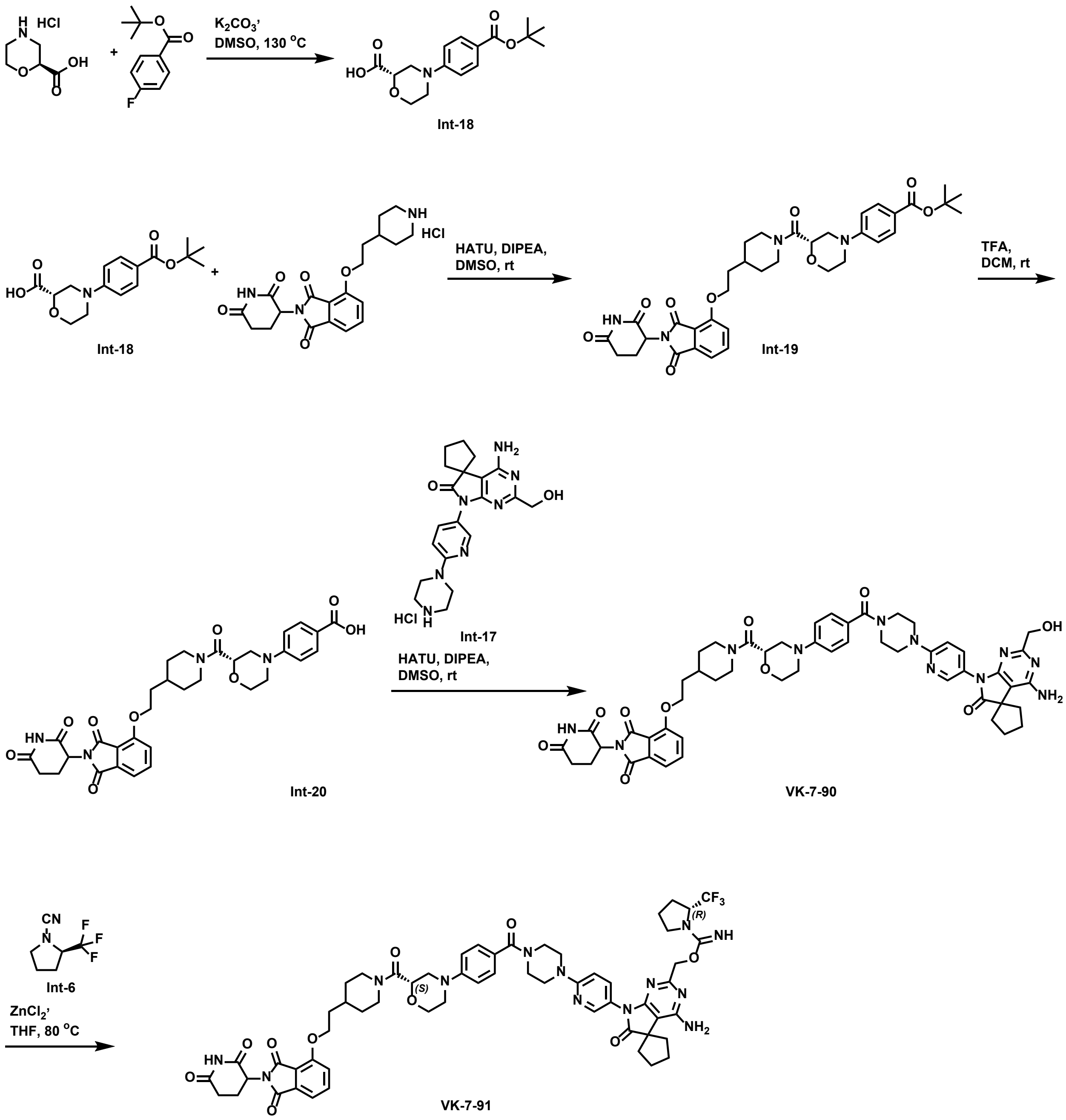


**(*S*)-4-(4-(tert-butoxycarbonyl)phenyl)morpholine-2-carboxylic acid (Int-18)**

To a solution of (*S*)-morpholine-2-carboxylic acid hydrochloride (300.0 mg, 2.29 mmol) and tert-butyl 4-fluorobenzoate (449.0 mg, 2.29 mmol) in DMSO (4.0 mL) was added $K_2CO_3$ (316.0 mg, 2.29 mmol) under $N_2$. The reaction mixture was stirred at 130 ºC for 18 h. After completion of the reaction, the mixture was cooled to rt, quenched with water (10.0 mL), and extracted with EtOAc (2 x 40.0 mL). The combined organic layers were dried over anhydrous $Na_2SO_4$, filtered, and concentrated under reduced pressure. The crude residue was purified via column chromatography (silica gel, eluted with 70% ethyl acetate in hexane) to give **Int-18** (310.0 mg, 56% yield) as a white solid. MS (ESI) for $C_{16}H_{21}NO_5$ $[M+H]^+$: m/z calcd, 308.15; found, 308.21.

**tert-butyl 4-((2*S*)-2-(4-(2-((2-(2,6-dioxopiperidin-3-yl)-1,3-dioxoisoindolin-4-yl)oxy)ethyl)piperidine-1-carbonyl)morpholino)benzoate (Int-19)**

To a solution of **Int-18** (150.0 mg, 0.49 mmol) and 2-(2,6-dioxopiperidin-3-yl)-4-(2-(piperidin-4-yl)ethoxy)isoindoline-1,3-dione hydrochloride (206.0 mg, 0.49 mmol, for the preparation method, see patent WO 2024003533 A1) in DMSO (1.2 mL) were added DIPEA (0.43 mL, 2.44 mmol) and HATU (223.0 mg, 0.59 mmol). The reaction mixture was stirred at rt for 30 minutes. The reaction mixture was purified directly via reverse-phase column chromatography on C-18 (eluted with 65% $CH_3CN$ in water) to afford **Int-19** (250.0 mg, 76% yield) as a white solid. MS (ESI) for $C_{36}H_{42}N_4O_9$ $[M+H]^+$: m/z calcd, 675.30; found, 675.40.

**4-((2*S*)-2-(4-(2-((2-(2,6-dioxopiperidin-3-yl)-1,3-dioxoisoindolin-4-yl)oxy)ethyl)piperidine-1-carbonyl)morpholino)benzoic acid (Int-20)**

To a solution of **Int-19** (40.0 mg, 59.28 μmol) in DCM (0.8 mL) was added TFA (0.2 mL) at 0 ºC. The reaction mixture was stirred at rt for 4 h. The reaction mixture was concentrated under reduced pressure to give **Int-20** (35.0 mg, 95% yield) as white solid. MS (ESI) for $C_{32}H_{34}N_4O_9$ $[M+H]^+$: m/z calcd, 619.24; found, 619.24.

**4-(2-(1-((*S*)-4-(4-(4-(5-(4'-amino-2'-(hydroxymethyl)-6'-oxospiro[cyclopentane-1,5'-pyrrolo[2,3-d]pyrimidin]-7'(6'H)-yl)pyridin-2-yl)piperazine-1-carbonyl)phenyl)morpholine-2-carbonyl)piperidin-4-yl)ethoxy)-2-(2,6-dioxopiperidin-3-yl)isoindoline-1,3-dione (VK-7-90)**

To a solution of **Int-17** (12.0 mg, 27.78 μmol) and **Int-20** (17.0 mg, 27.78 μmol) in DMSO (0.5 mL) were added DIPEA (24.0 μL, 0.14 mmol) and HATU (12.7 mg, 33.33 μmol). The reaction mixture was stirred at rt for 30 minutes. The reaction mixture was purified directly via reverse-phase column chromatography on C-18 (eluted with 45% $CH_3CN$ in water) to afford **VK-7-90** (18.0 mg, 65% yield) as a white solid. $^1$H NMR (600 MHz, DMSO-$d_6$) δ 11.09 (s, 1H), 8.14 (d, *J* = 2.7 Hz, 1H), 7.81 (dd, *J* = 8.5, 7.3 Hz, 1H), 7.59 (dd, *J* = 9.1, 2.7 Hz, 1H), 7.54 (d, *J* = 8.5 Hz, 1H), 7.45 (d, *J* = 7.3 Hz, 1H), 7.37 (d, *J* = 8.8 Hz, 2H), 7.00 (d, *J* = 8.8 Hz, 2H), 6.94 (d, *J* = 9.1 Hz, 1H), 6.52 (brs, 2H), 5.08 (dd, *J* = 13.0, 5.6 Hz, 1H), 4.75 (t, *J* = 5.9 Hz, 1H), 4.41-4.31 (m, 2H), 4.30-4.25 (m, 2H), 4.23 (d, *J* = 5.9 Hz, 2H), 4.03-3.94 (m, 2H), 3.79-3.72 (m, 1H), 3.70-3.58 (m, 10H), 3.11-2.75 (m, 4H), 2.63-2.52 (m, 2H), 2.18-2.11 (m, 2H), 2.04-1.94 (m, 5H), 1.92-1.67 (m, 8H), 1.28-1.02 (m, 2H). MS (ESI) for $C_{52}H_{57}N_{11}O_{10}$ $[M+H]^+$: m/z calcd, 996.44; found, 996.46.

**(4'-amino-7'-(6-(4-(4-((2*S*)-2-(4-(2-((2-(2,6-dioxopiperidin-3-yl)-1,3-dioxoisoindolin-4-yl)oxy)ethyl)piperidine-1-carbonyl)morpholino)benzoyl)piperazin-1-yl)pyridin-3-yl)-6'-oxo-6',7'-dihydrospiro[cyclopentane-1,5'-pyrrolo[2,3-d]pyrimidin]-2'-yl)methyl (2*R*)-2-(trifluoromethyl)pyrrolidine-1-carbimidate (VK-7-91)**

Following the same procedure as compound **VK-4-119** by using **VK-7-90** instead of **Int-12**, compound **VK-7-91** (12.0 mg, 57% yield) was prepared as a white solid. $^1$H NMR (600 MHz, DMSO-$d_6$) δ 11.08 (s, 1H), 8.12 (d, *J* = 2.6 Hz, 1H), 7.81 (dd, *J* = 8.6, 7.3 Hz, 1H), 7.58 (dd, *J* = 9.1, 2.6 Hz, 1H), 7.54 (d, *J* = 8.6 Hz, 1H), 7.45 (d, *J* = 7.3 Hz, 1H), 7.37 (d, *J* = 8.4 Hz, 2H), 7.00 (d, *J* = 8.2 Hz, 2H), 6.91 (d, *J* = 9.1 Hz, 1H), 6.53 (brs, 2H), 5.75 (s, 1H), 5.08 (dd, *J* = 12.8, 5.5

Hz, 1H), 5.02-4.78 (m, 2H), 4.56-4.44 (m, 1H), 4.40-4.31 (m, 2H), 4.30-4.24 (m, 2H), 4.04-3.93 (m, 2H), 3.78-3.72 (m, 1H), 3.70-3.56 (m, 10H), 3.27-3.21 (m, 1H), 3.08-2.93 (m, 2H), 2.91-2.79 (m, 2H), 2.67-2.51 (m, 3H), 2.18-2.10 (m, 2H), 2.05-1.94 (m, 5H), 1.94-1.67 (m, 12H), 1.28-1.03 (m, 2H). MS (ESI) for $C_{58}H_{64}F_3N_{13}O_{10}$ $[M+H]^+$: m/z calcd, 1160.49; found, 1160.58.

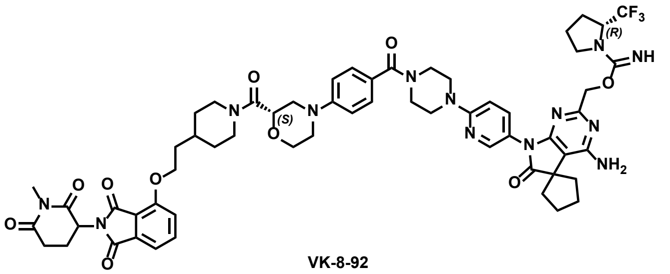

VK-8-92

**(4'-amino-7'-(6-(4-(4-((2*S*)-2-(4-(2-((2-(1-methyl-2,6-dioxopiperidin-3-yl)-1,3-dioxoisoindolin-4-yl)oxy)ethyl)piperidine-1-carbonyl)morpholino)benzoyl)piperazin-1-yl)pyridin-3-yl)-6'-oxo-6',7'-dihydrospiro[cyclopentane-1,5'-pyrrolo[2,3-d]pyrimidin]-2'-yl)methyl (2*R*)-2-(trifluoromethyl)pyrrolidine-1-carbimidate (VK-8-92)**

Following the same procedure as compound **VK-7-91** by using 4-(2-(1-((*S*)-4-(4-(4-(5-(4'-amino-2'-(hydroxymethyl)-6'-oxospiro[cyclopentane-1,5'-pyrrolo[2,3-d]pyrimidin]-7'(6'H)-yl)pyridin-2-yl)piperazine-1-carbonyl)phenyl)morpholine-2-carbonyl)piperidin-4-yl)ethoxy)-2-(1-methyl-2,6-dioxopiperidin-3-yl)isoindoline-1,3-dione instead of **VK-7-90**, compound **VK-8-92** (8.0 mg, 69% yield) was prepared as a white solid. $^1$H NMR (600 MHz, DMSO-$d_6$) δ 8.11 (d, $J$ = 2.6 Hz, 1H), 7.82 (dd, $J$ = 8.5, 7.2 Hz, 1H), 7.57 (dd, $J$ = 9.1, 2.6 Hz, 1H), 7.54 (d, $J$ = 8.5 Hz, 1H), 7.45 (d, $J$ = 7.2 Hz, 1H), 7.37 (d, $J$ = 8.8 Hz, 2H), 7.00 (d, $J$ = 8.8 Hz, 2H), 6.91 (d, $J$ = 9.1 Hz, 1H), 6.57 (brs, 2H), 5.15 (dd, $J$ = 12.7, 5.4 Hz, 1H), 5.07-4.87 (m, 2H), 4.60-4.50 (m, 1H), 4.40-4.24 (m, 4H), 4.04-3.93 (m, 2H), 3.78-3.72 (m, 1H), 3.70-3.57 (m, 10H), 3.31-3.23 (m, 1H), 3.08-3.00 (m, 1H), 3.01 (s, 3H), 2.98-2.90 (m, 2H), 2.85-2.73 (m, 2H), 2.65-2.51 (m, 2H), 2.18-2.11 (m, 2H), 2.07-1.70 (m, 17H), 1.27-1.03 (m, 2H). MS (ESI) for $C_{59}H_{66}F_3N_{13}O_{10}$ $[M+H]^+$: m/z calcd, 1174.51; found, 1174.48. The exchangeable proton signal of the -C=NH group was not observed in $^1$H NMR. This also applies to the following other degrader compounds.

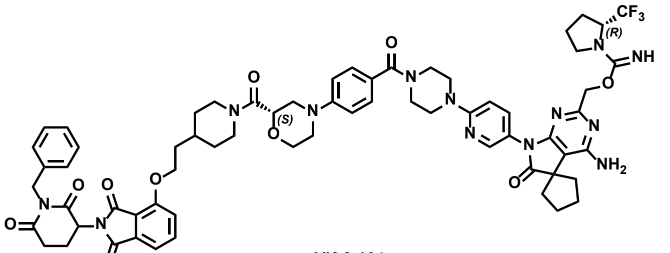

VK-8-164

**(4'-amino-7'-(6-(4-(4-((2*S*)-2-(4-(2-((2-(1-benzyl-2,6-dioxopiperidin-3-yl)-1,3-dioxoisoindolin-4-yl)oxy)ethyl)piperidine-1-carbonyl)morpholino)benzoyl)piperazin-1-yl)pyridin-3-yl)-6'-oxo-6',7'-dihydrospiro[cyclopentane-1,5'-pyrrolo[2,3-d]pyrimidin]-2'-yl)methyl (2*R*)-2-(trifluoromethyl)pyrrolidine-1-carbimidate (VK-8-164)**

Following the same procedure as compound **VK-7-91** by using 4-(2-(1-((S)-4-(4-(4-(5-(4'-amino-2'-(hydroxymethyl)-6'-oxospiro[cyclopentane-1,5'-pyrrolo[2,3-d]pyrimidin]-7'(6'H)-yl)pyridin-2-yl)piperazine-1-carbonyl)phenyl)morpholine-2-carbonyl)piperidin-4-yl)ethoxy)-2-(1-benzyl-2,6-dioxopiperidin-3-yl)isoindoline-1,3-dione instead of **VK-7-90**, compound **VK-8-164** (8.5 mg, 41% yield) was prepared as a white solid. $^1$H NMR (600 MHz, DMSO-$d_6$) δ 8.12 (d, $J$ = 2.6 Hz, 1H), 7.82 (dd, $J$ = 8.5, 7.3 Hz, 1H), 7.57 (dd, $J$ = 9.1, 2.6 Hz, 1H), 7.55 (d, $J$ = 8.5 Hz, 1H), 7.46 (d, $J$ = 7.3 Hz, 1H), 7.37 (d, $J$ = 8.6 Hz, 2H), 7.32-7.28 (m, 2H), 7.26-7.21 (m, 3H), 7.00 (dd, $J$ = 9.1, 2.7 Hz, 2H), 6.92 (d, $J$ = 9.1 Hz, 1H), 6.56 (brs, 2H), 5.33-5.28 (m, 1H), 4.98-4.77 (m, 4H), 4.54-4.46 (m, 1H), 4.40-4.25 (m, 4H), 4.03-3.94 (m, 2H), 3.72-3.77 (m, 1H), 3.70-3.56 (m, 10H), 3.29-3.26 (m, 1H), 3.11-2.92 (m, 3H), 2.84-2.77 (m, 2H), 2.65-2.55 (m, 2H), 2.17-2.06 (m, 3H), 2.00-1.94 (m, 4H), 1.94-1.68 (m, 12H), 1.28-1.03 (m, 2H). MS (ESI) for $C_{65}H_{70}F_3N_{13}O_{10}$ $[M/2+H]^+$: m/z calcd, 625.76; found, 625.87.

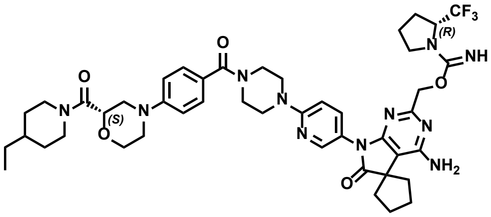


VK-8-129

**(4'-amino-7'-(6-(4-(4-((*S*)-2-(4-ethylpiperidine-1-carbonyl)morpholino)benzoyl)piperazin-1-yl)pyridin-3-yl)-6'-oxo-6',7'-dihydrospiro[cyclopentane-1,5'-pyrrolo[2,3-d]pyrimidin]-2'-yl)methyl (*R*)-2-(trifluoromethyl)pyrrolidine-1-carbimidate (VK-8-129)**

Following the same procedure as compound **VK-7-91** by using (*S*)-4'-amino-7'-(6-(4-(4-(2-(4-ethylpiperidine-1-carbonyl)morpholino)benzoyl)piperazin-1-yl)pyridin-3-yl)-2'-(hydroxymethyl)spiro[cyclopentane-1,5'-pyrrolo[2,3-d]pyrimidin]-6'(7'H)-one instead of **VK-7-90**, compound **VK-8-129** (9.0 mg, 67% yield) was prepared as a white solid. $^1$H NMR (600 MHz, DMSO-$d_6$) δ 8.11 (d, $J$ = 2.6 Hz, 1H), 7.57 (dd, $J$ = 9.1, 2.6 Hz, 1H), 7.37 (d, $J$ = 8.8 Hz, 2H), 7.00 (dd, $J$ = 9.1, 2.6 Hz, 2H), 6.92 (d, $J$ = 9.1 Hz, 1H), 6.57 (brs, 2H), 5.04-4.84 (m, 2H), 4.57-4.48 (m, 1H), 4.40-4.32 (m, 2H), 4.03-3.94 (m, 2H), 3.78-3.72 (m, 1H), 3.70-3.55 (m, 10H), 3.31-3.25 (m, 2H), 3.04-2.78 (m, 3H), 2.62-2.52 (m, 1H), 2.18-2.10 (m, 2H), 2.00-1.67 (m, 12H), 1.43-1.36 (m, 1H), 1.27-1.21 (m, 2H), 1.13-0.89 (m, 2H), 0.87 (t, $J$ = 7.3 Hz, 3H). MS (ESI) for $C_{45}H_{56}F_3N_{11}O_5$ $[M+H]^+$: m/z calcd, 888.45; found, 888.55.

### NMR spectra of key compounds

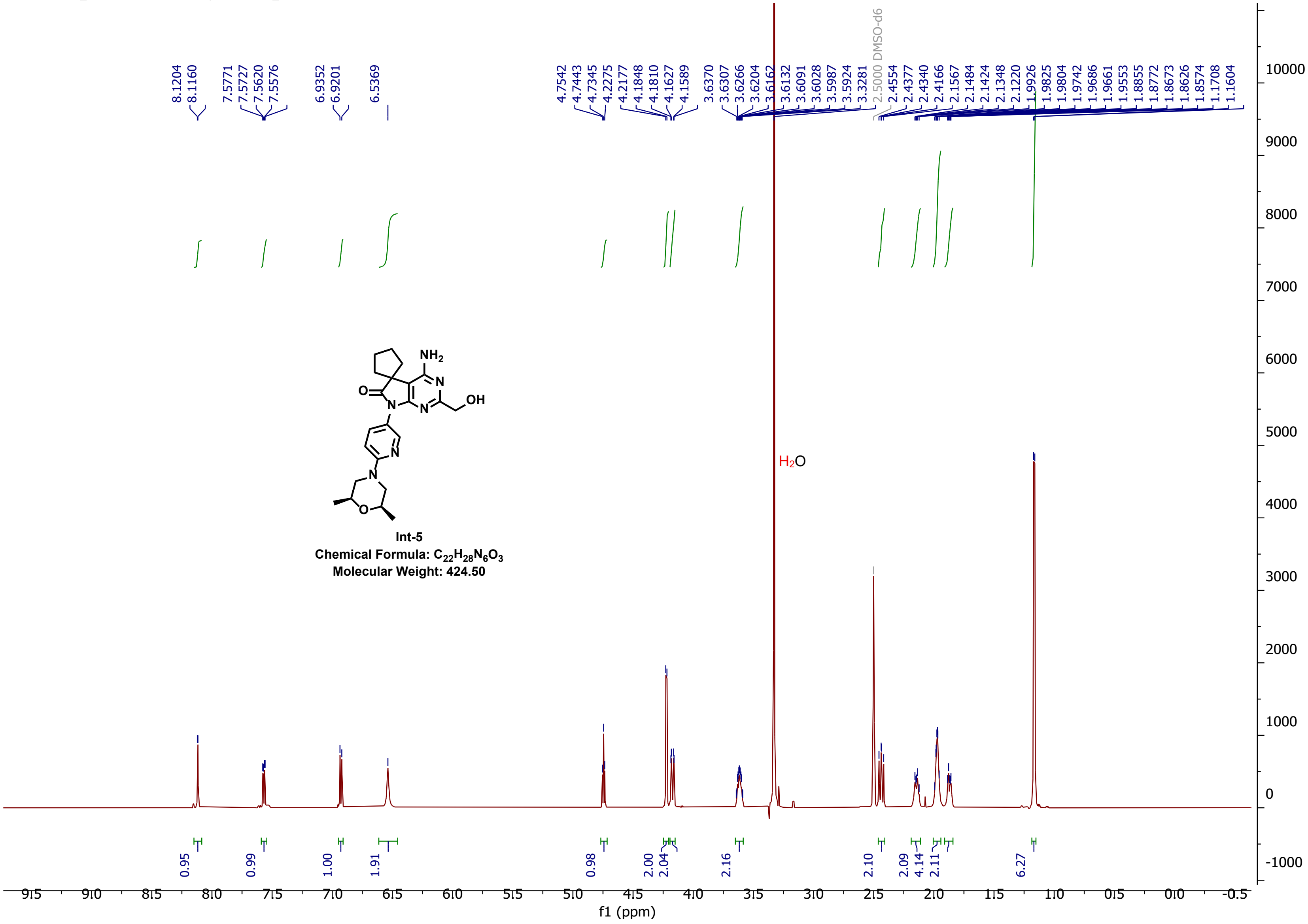

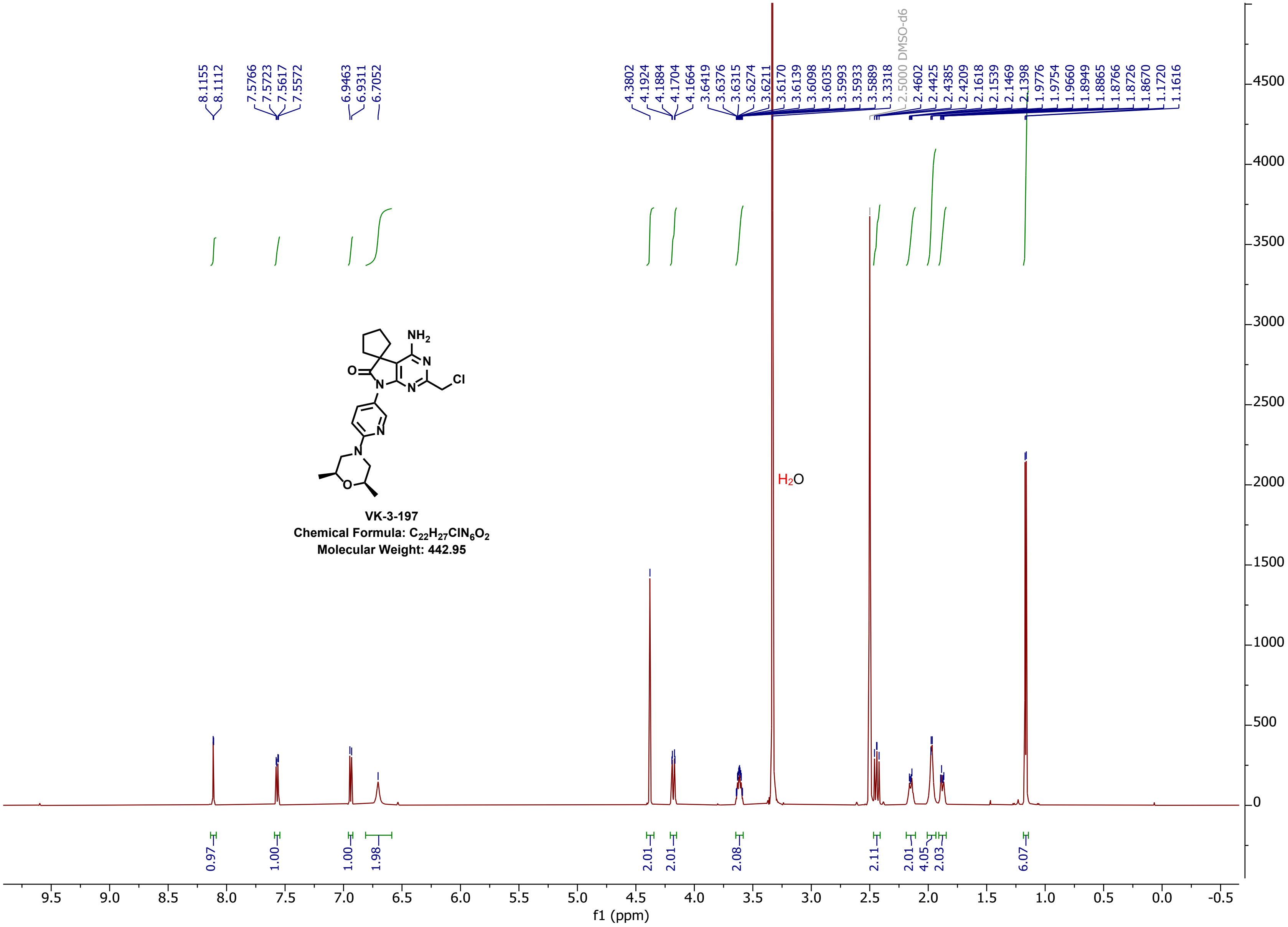

VK-3-197
Chemical Formula: $C_{22}H_{27}ClN_6O_2$
Molecular Weight: 442.95

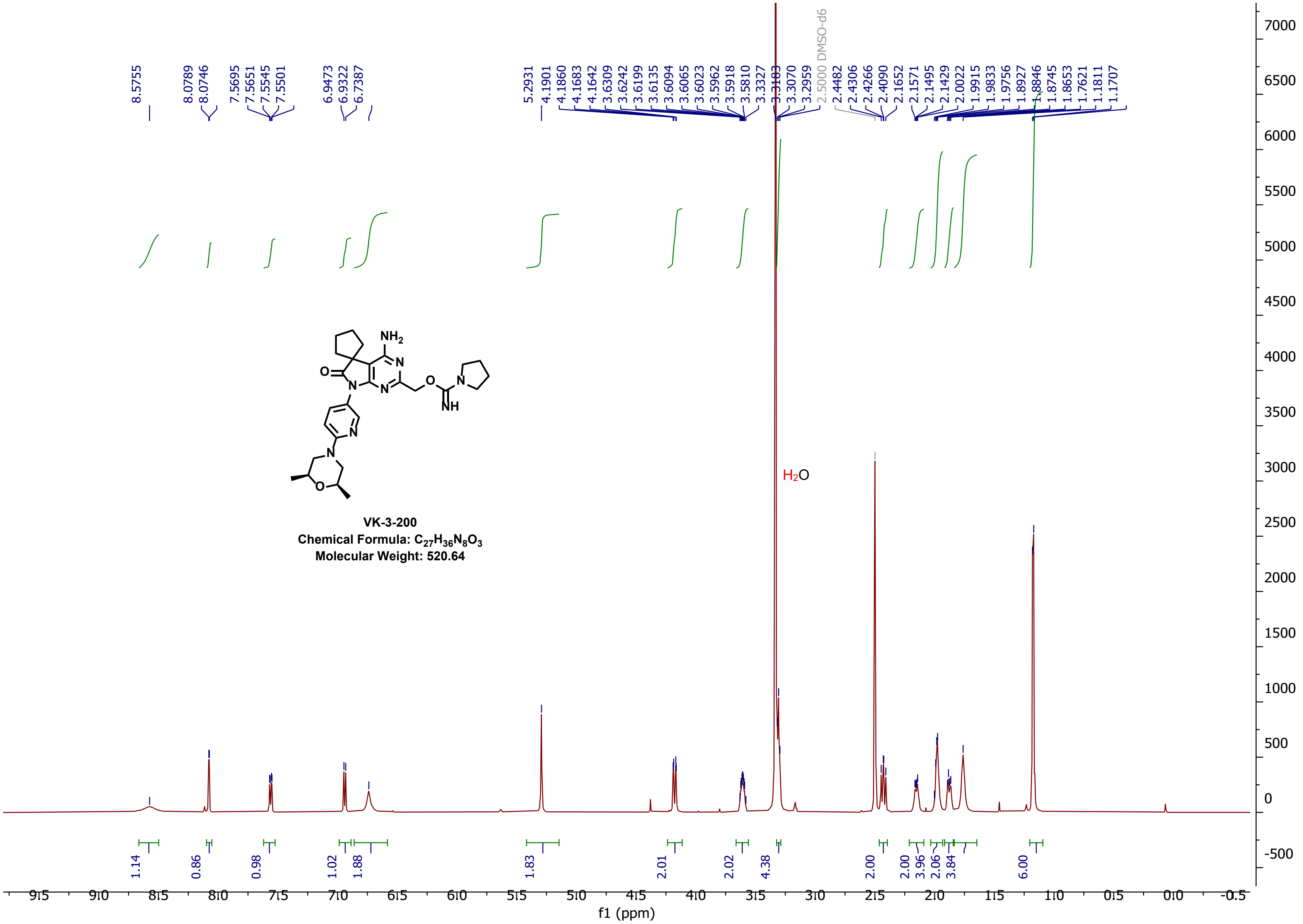

8.5755
8.0789
8.0746
7.5695
7.5651
7.5545
7.5501
6.9473
6.9322
6.7387
5.2931
4.1901
4.1860
4.1683
4.1642
3.6309
3.6242
3.6199
3.6135
3.6094
3.6065
3.6023
3.5962
3.5918
3.5810
3.3327
3.3183
3.3070
3.2959
2.5000 DMSO-d6
2.4482
2.4306
2.4266
2.4090
2.1652
2.1571
2.1495
2.1429
2.0022
1.9915
1.9833
1.9756
1.8927
1.8846
1.8745
1.8653
1.7621
1.1811
1.1707
H2O
NH2
N
O
NH
VK-3-200
Chemical Formula: C27H36N8O3
Molecular Weight: 520.64
1.14
0.86
0.98
1.02
1.88
1.83
2.01
2.02
4.38
2.00
3.96
2.06
3.84
6.00
f1 (ppm)

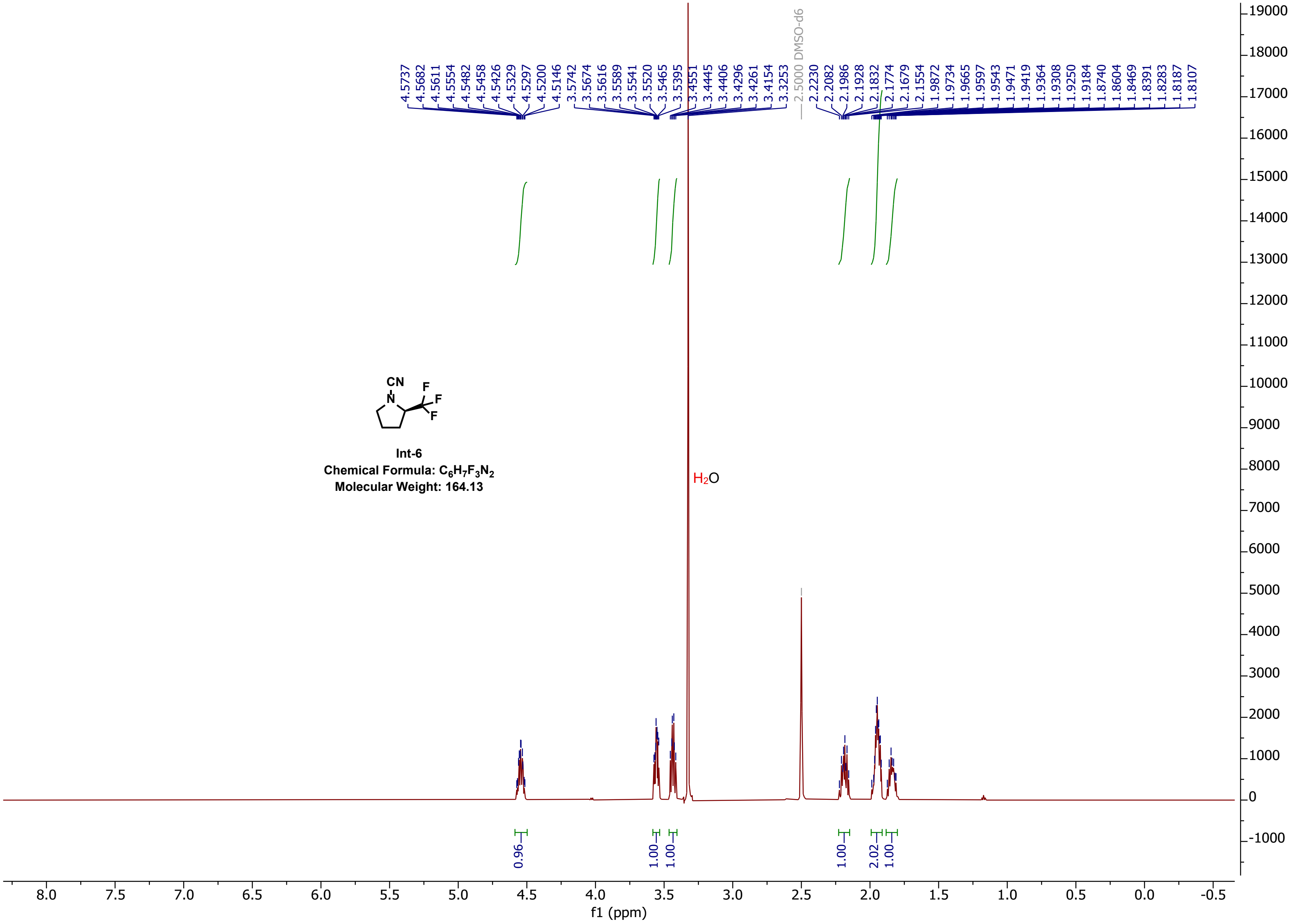
4.5737
4.5682
4.5611
4.5554
4.5482
4.5458
4.5426
4.5329
4.5297
4.5200
4.5146
3.5742
3.5674
3.5616
3.5589
3.5541
3.5520
3.5465
3.5395
3.4551
3.4445
3.4406
3.4296
3.4261
3.4154
3.3253
2.5000 DMSO-d6
2.2230
2.2082
2.1986
2.1928
2.1832
2.1774
2.1679
2.1554
1.9872
1.9734
1.9665
1.9597
1.9543
1.9471
1.9419
1.9364
1.9308
1.9250
1.9184
1.8740
1.8604
1.8469
1.8391
1.8283
1.8187
1.8107
CN
N
F
F
F
Int-6
Chemical Formula: $C_6H_7F_3N_2$
Molecular Weight: 164.13
$H_2O$
0.96
1.00
1.00
1.00
2.02
1.00
f1 (ppm)

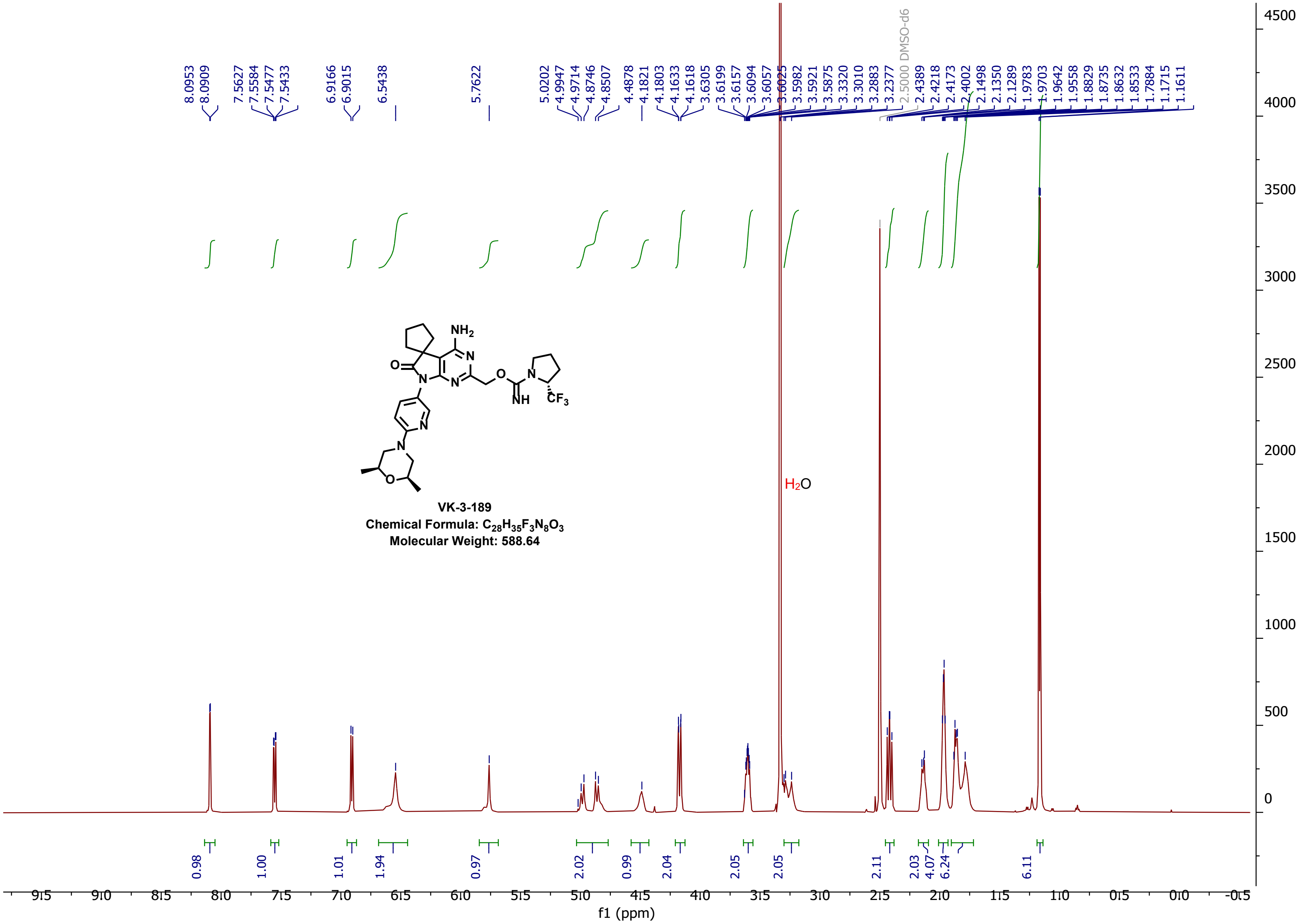
8.0953
8.0909
7.5627
7.5584
7.5477
7.5433
6.9166
6.9015
6.5438
5.7622
5.0202
4.9947
4.9714
4.8746
4.8507
4.4878
4.1821
4.1803
4.1633
4.1618
3.6305
3.6199
3.6157
3.6094
3.6057
3.6025
3.5982
3.5921
3.5875
3.3320
3.3010
3.2883
3.2377
2.5000 DMSO-d6
2.4389
2.4218
2.4173
2.4002
2.1498
2.1350
2.1289
1.9783
1.9703
1.9642
1.9558
1.8829
1.8735
1.8632
1.8533
1.7884
1.1715
1.1611
NH2
N
O
N
N
O
N
NH
CF3
N
N
O
VK-3-189
Chemical Formula: C28H35F3N8O3
Molecular Weight: 588.64
H2O
0.98
1.00
1.01
1.94
0.97
2.02
0.99
2.04
2.05
2.05
2.11
2.03
4.07
6.24
6.11
f1 (ppm)

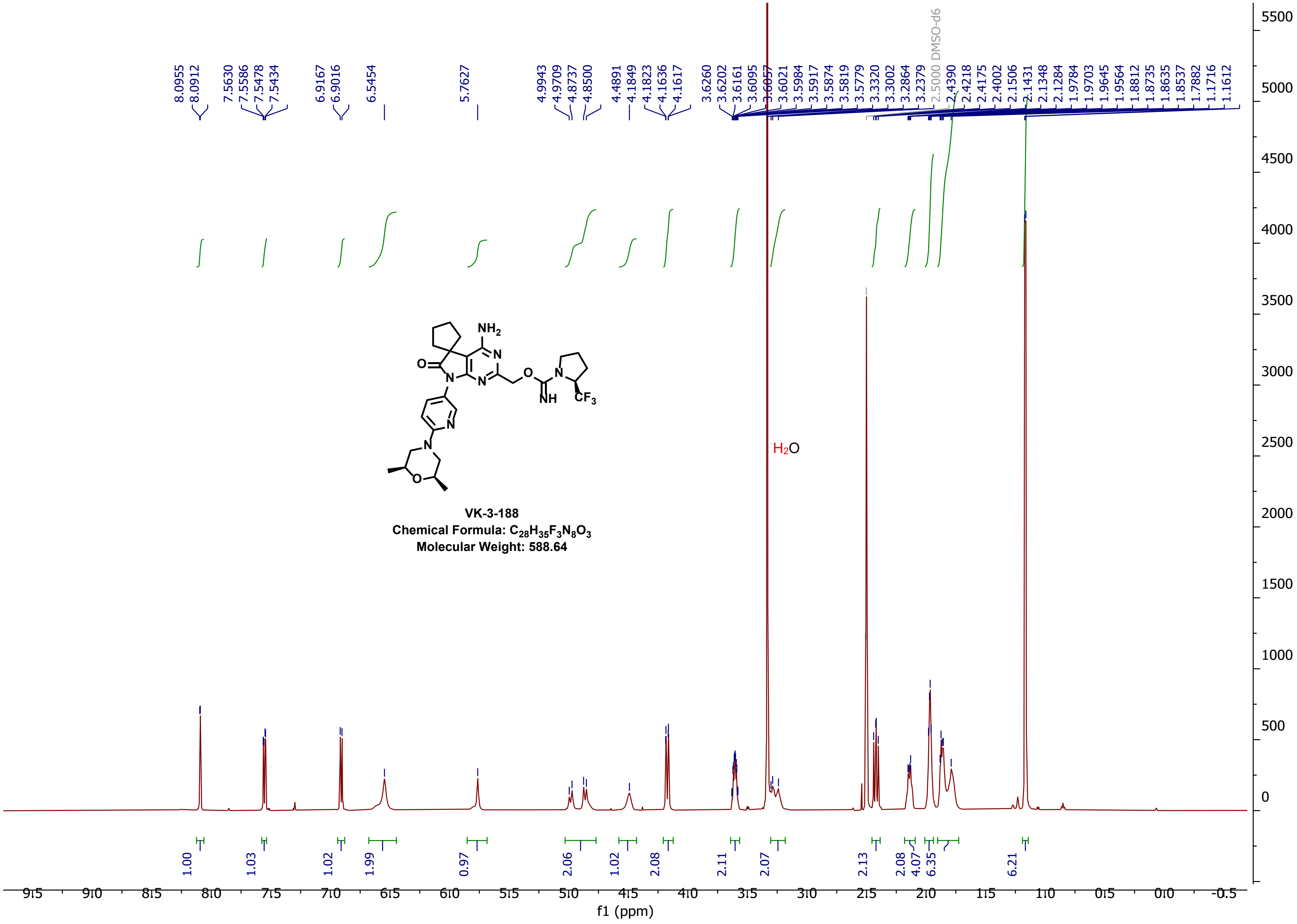
8.0955
8.0912
7.5630
7.5586
7.5478
7.5434
6.9167
6.9016
6.5454
5.7627
4.9943
4.9709
4.8737
4.8500
4.4891
4.1849
4.1823
4.1636
4.1617
3.6260
3.6202
3.6161
3.6095
3.6057
3.6021
3.5984
3.5917
3.5874
3.5819
3.5779
3.3320
3.3002
3.2864
3.2379
2.5000 DMSO-d6
2.4390
2.4218
2.4175
2.4002
2.1506
2.1431
2.1348
2.1284
1.9784
1.9703
1.9645
1.9564
1.8812
1.8735
1.8635
1.8537
1.7882
1.1716
1.1612
NH2
O
N
N
N
O
N
NH
CF3
N
N
O
VK-3-188
Chemical Formula: C28H35F3N8O3
Molecular Weight: 588.64
H2O
1.00
1.03
1.02
1.99
0.97
2.06
1.02
2.08
2.11
2.07
2.13
2.08
4.07
6.35
6.21
f1 (ppm)

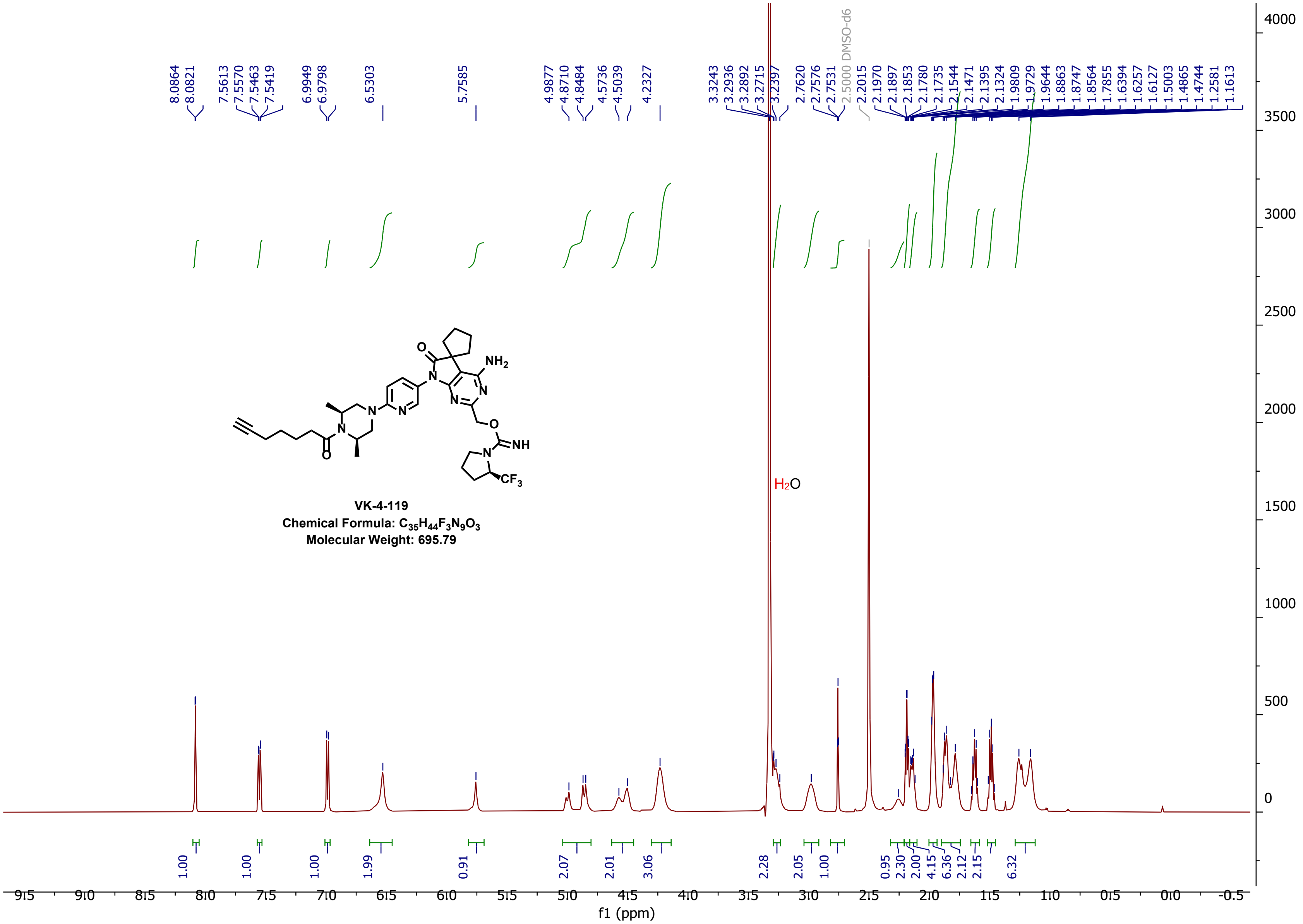
VK-4-119
Chemical Formula: $C_{35}H_{44}F_3N_9O_3$
Molecular Weight: 695.79
$H_2O$
2.5000 DMSO-d6
f1 (ppm)

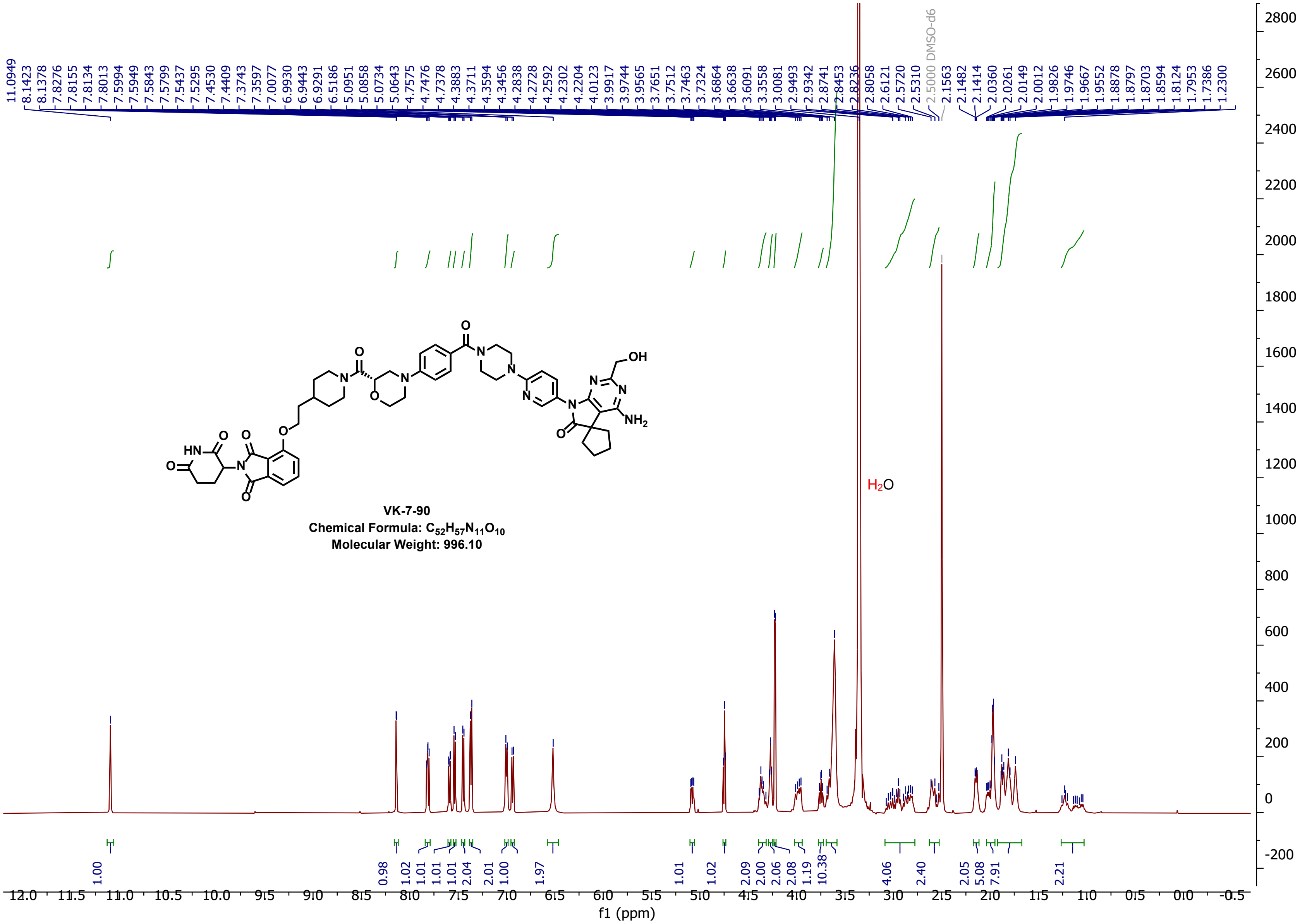

**VK-7-90**
**Chemical Formula: $C_{52}H_{57}N_{11}O_{10}$**
**Molecular Weight: 996.10**

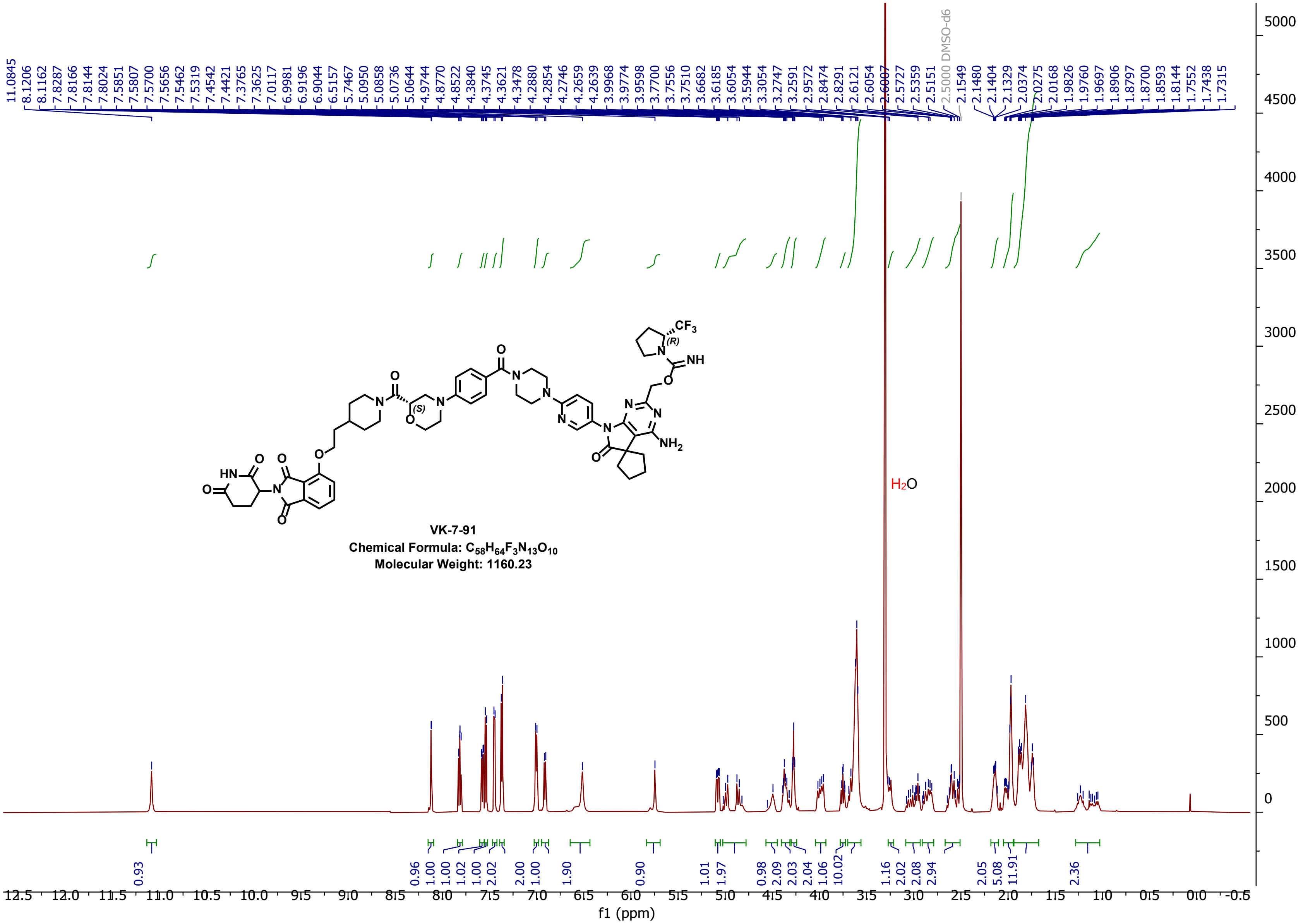
11.0845
8.1206
8.1162
7.8287
7.8166
7.8144
7.8024
7.5851
7.5807
7.5700
7.5656
7.5462
7.5319
7.4542
7.4421
7.3765
7.3625
7.0117
6.9981
6.9196
6.9044
6.5157
5.7467
5.0950
5.0858
5.0736
5.0644
4.9744
4.8770
4.8522
4.3840
4.3745
4.3621
4.3478
4.2880
4.2854
4.2746
4.2659
4.2639
3.9968
3.9774
3.9598
3.7700
3.7556
3.7510
3.6682
3.6185
3.6054
3.5944
3.3054
3.2747
3.2591
2.9572
2.8474
2.8291
2.6121
2.6054
2.6007
2.5727
2.5359
2.5151
2.5000 DMSO-d6
2.1549
2.1480
2.1404
2.1329
2.0374
2.0275
2.0168
1.9826
1.9760
1.9697
1.8906
1.8797
1.8700
1.8593
1.8144
1.7552
1.7438
1.7315
H2O
VK-7-91
Chemical Formula: C58H64F3N13O10
Molecular Weight: 1160.23
0.93
0.96
1.00
1.00
1.02
1.00
2.02
2.00
1.00
1.90
0.90
1.01
1.97
0.98
2.09
2.03
2.04
1.06
10.02
1.16
2.02
2.08
2.94
2.05
5.08
11.91
2.36
f1 (ppm)

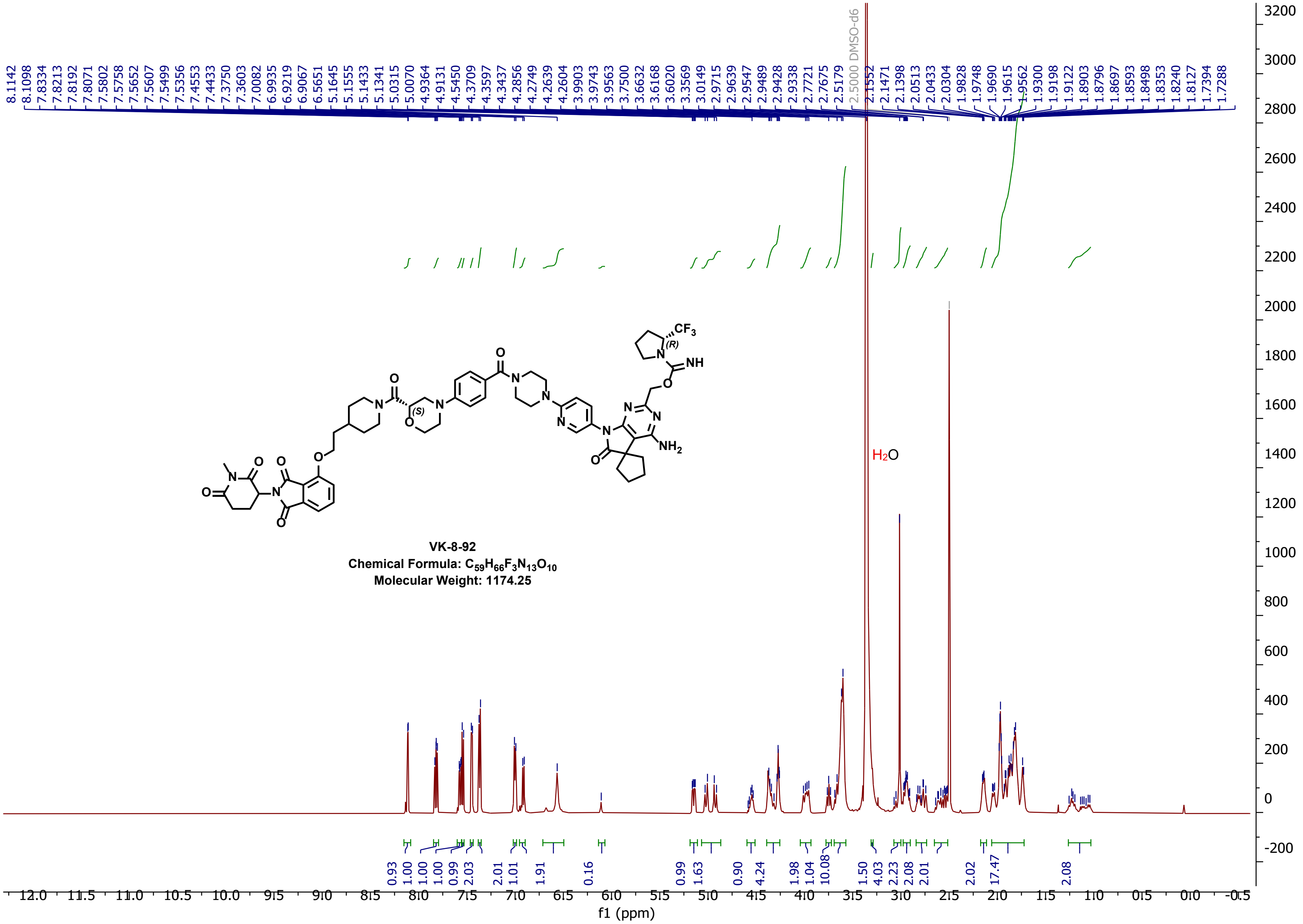
VK-8-92
Chemical Formula: $C_{59}H_{66}F_3N_{13}O_{10}$
Molecular Weight: 1174.25
DMSO-d6
$H_2O$
f1 (ppm)

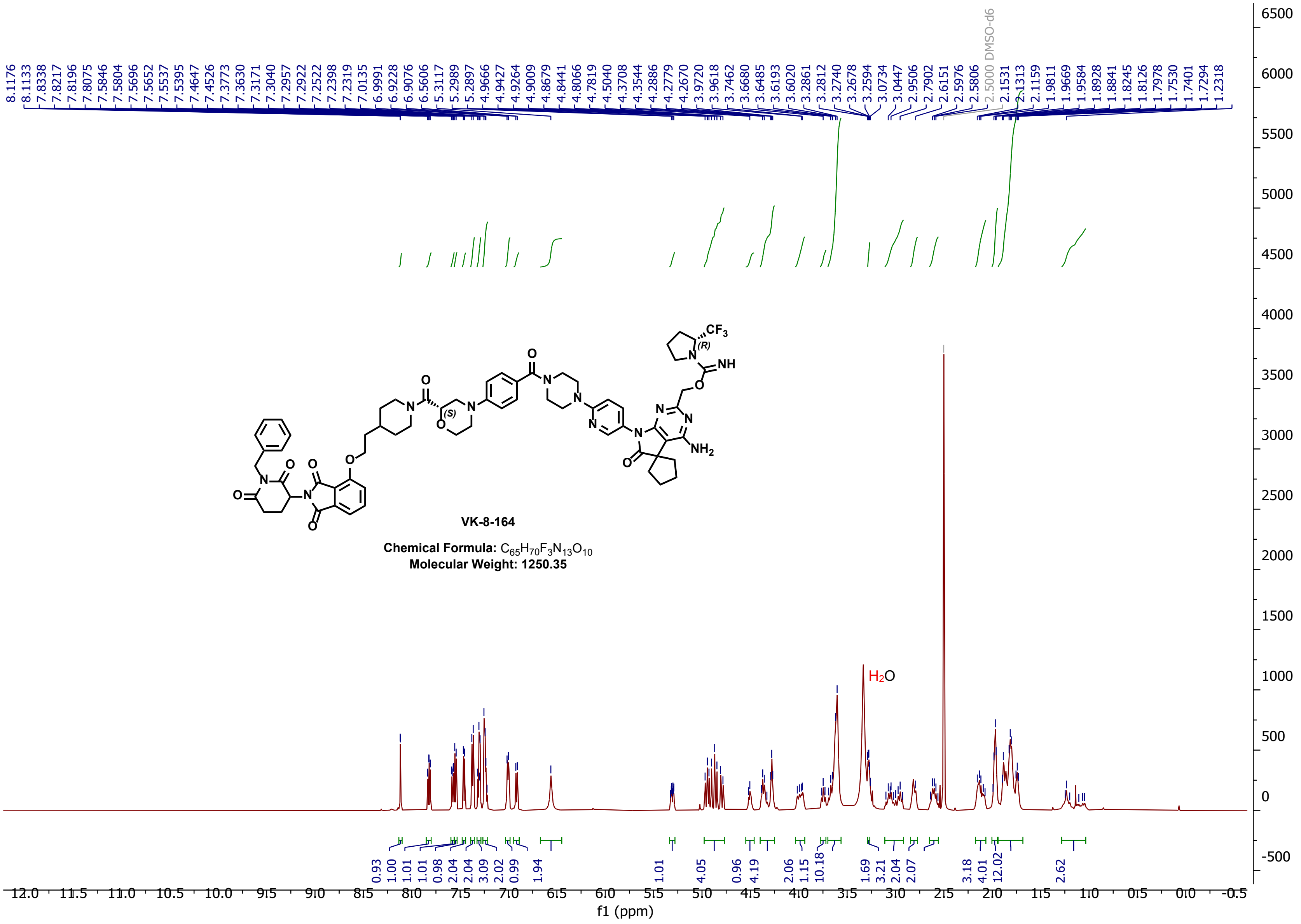

VK-8-164
Chemical Formula: C65H70F3N13O10
Molecular Weight: 1250.35
H2O
2.5000 DMSO-d6
f1 (ppm)

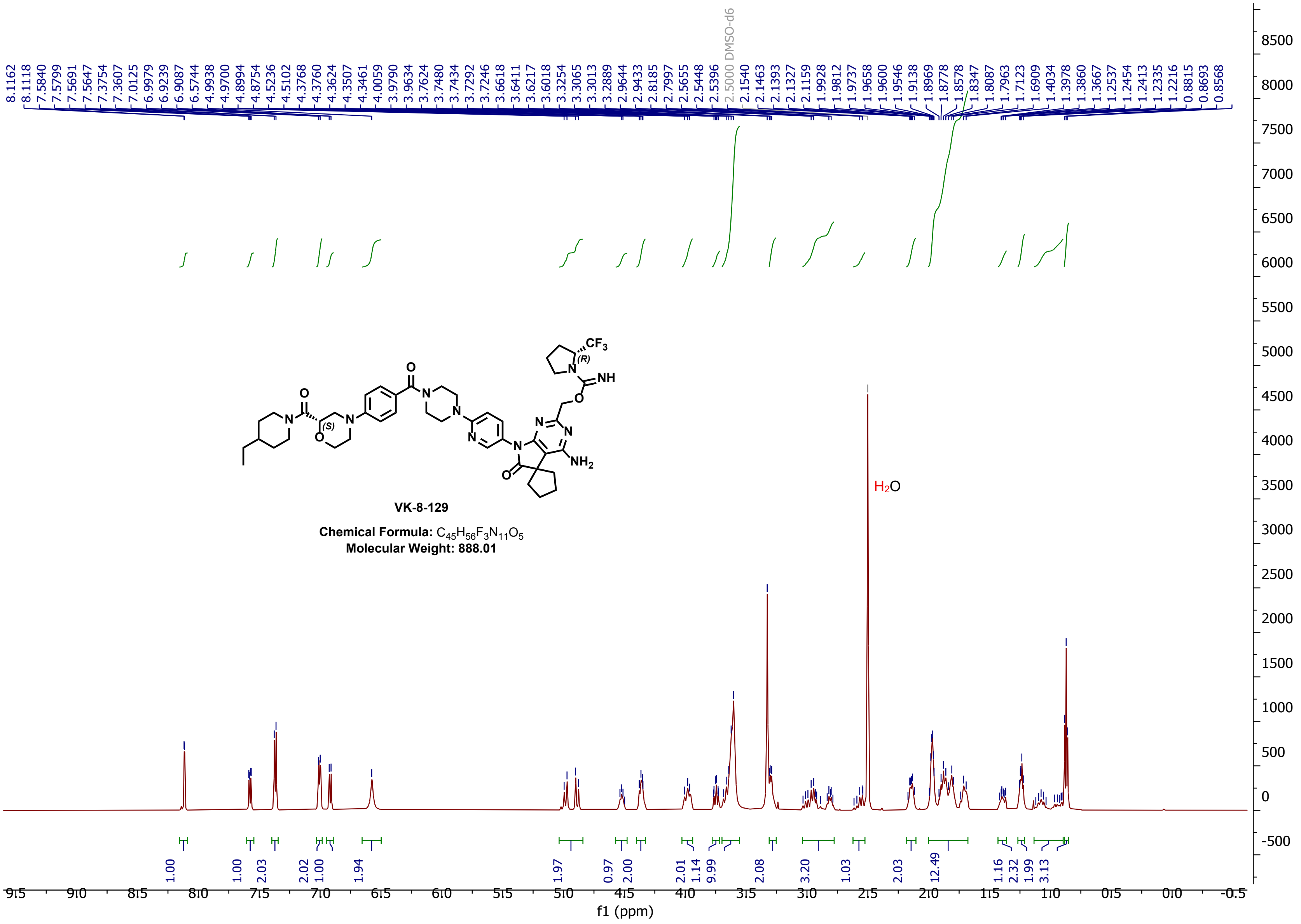

DMSO-d6
VK-8-129
Chemical Formula: C45H56F3N11O5
Molecular Weight: 888.01
H2O
f1 (ppm)
